# Integrative analysis of patient-derived tumoroids and ex vivo organoid modeling of ARID1A loss in bladder cancer reveals therapeutic molecular targets

**DOI:** 10.1101/2024.07.29.601702

**Authors:** Mathijs P. Scholtes, Maryam Akbarzadeh, J. Alberto Nakauma-Gonzáles, Alexandros Galaras, Ameneh Bazrafshan, Bram Torenvliet, Leila Beikmohammadi, Valeria Lozovanu, Shahla Romal, Panagiotis Moulos, Tsung Wai Kan, Mahesh Algoe, Martin E. van Royen, Andrea Sachetti, Thierry P.P. van den Bosch, Bert Eussen, Annelies de Klein, Geert J.L.H. van Leenders, Joost L. Boormans, Pantelis Hatzis, Robert-Jan Palstra, Tahlita C.M. Zuiverloon, Tokameh Mahmoudi

## Abstract

Somatic mutations in *ARID1A* (AT-rich interactive domain-containing protein 1A) are present in approximately 25% of bladder cancers (BC) and are associated with poor prognosis. With a view to discover effective treatment options for ARID1A-deficient BC patients, we set out to identify targetable effectors dysregulated consequent to ARID1A deficiency. Integrative analyses of ARID1A depletion in normal organoids and data mining in publicly available datasets revealed upregulation of DNA repair and cell cycle-associated genes consequent to loss of ARID1A and identified *CHEK1* (Checkpoint kinase 1) and chromosomal passenger complex member *BIRC5* (Baculoviral IAP Repeat Containing 5) as therapeutically drug-able candidate molecular effectors. *Ex vivo* treatment of patient-derived BC tumoroids with clinically advanced small molecule inhibitors targeting *CHEK1* or *BIRC5* was associated with increased DNA damage signalling and apoptosis, and selectively induced cell death in tumoroids lacking ARID1A protein expression. Thus, integrating public datasets with patient-derived organoid modelling and ex-vivo drug testing can uncover key molecular effectors and mechanisms of oncogenic transformation, potentially leading to novel therapeutic strategies. Our data point to ARID1A protein expression as a suitable candidate biomarker for the selection of BC patients responsive to therapies targeting BIRC5 and CHEK1.

## INTRODUCTION

Bladder cancer (BC) represents a significant global health burden, ranking as the 12th most prevalent malignancy worldwide and accounting for approximately 200,000 annual deaths[1]. Non-muscle invasive BC (NMIBC) patients have a favourable prognosis and receive local treatment. Management of muscle-invasive BC (MIBC), however, remains challenging due to its propensity for metastasis[2]. Standard-of-care in non-metastatic MIBC patients is cisplatin-based pre-operative chemotherapy and a radical cystectomy (RC)[2]. Despite the toxic side effects of pre-operative chemotherapy, half of MIBC patients will progress to metastatic urothelial cancer (mUC), which is characterized by a 5-year survival rate of approximately 5% [2, 3]. Poor mUC survival outcome highlights the clinical need for additional treatment options.

BC represents an example of how chromatin misregulation leads to cancer[4]. More than 80% of patients with BC harbour cancer-associated mutations in chromatin remodelling genes[4]. One of the most frequently mutated chromatin remodellers in MIBC and mUC is the AT-rich interactive domain-containing protein 1A (*ARID1A*), the defining component of the BAF ATP-dependent SWI/SNF nucleosome remodelling complex[4, 5]. Somatic mutations in *ARID1A* are present in approximately 20-30% of cases of MIBC and mUC, and predominantly include nonsense, point, and insertion or deletion frameshift mutations[4, 6, 7]. These mutations typically result in truncated proteins or reduced protein expression[8]. *ARID1A* is classified as a tumor suppressor because its genetic deletion impairs DNA double-strand break repair, disrupts telomere cohesion, and results in the upregulation of oncogenes[9–12]. The role of *ARID1A* as a global chromatin conformation regulator underlies the diverse effects observed when this gene is disrupted. Cellular processes impaired by loss of *ARID1A* serve as therapeutic targets for *ARID1A*-deficient BC[8, 11, 12]. Pharmacological inhibition of such targets could enable therapies which both exploit tumor-specific gene alterations and reduce overall toxicity.

Although many novel targets are pre-clinically identified for BC, few therapies are implemented in clinical practice. A major bottleneck is the lack of patient-representative preclinical models for candidate drug discovery and validation of clinically effective treatments[13, 14]. Recently, patient-derived tumoroids (malignant) and organoids (non-malignant) have been shown to be robust *ex vivo* platforms that recapitulate many attributes of human tissues, including 3-dimensional structure, multilineage differentiation, histological features, functional characteristics, and patient-treatment responses[15–20]. Additionally, their capacity for genetic manipulation makes tumoroids and organoids an ideal platform for pre-clinical BC research[18–21].

In this study, we explored novel pre-clinical treatment options for *ARID1A*-deficient BC. Our analysis of publicly available sequencing data from MIBC and mUC revealed that genetic aberrations in *ARID1A* are associated with reduced gene expression and poor patient prognosis. Next, we established and characterized a patient-derived BC tumoroid and normal urothelial organoid biobank and utilized this platform to identify dysregulated cellular processes and therapeutic candidate genes consequent to *ARID1A* deficiency. Finally, we investigated *BIRC5* and *CHEK1* as potential pharmacological targets to selectively eliminate ARID1A-deficient BC tumoroids.

## RESULTS

### *ARID1A* mutations correlate with reduced *ARID1A* gene expression and poor outcome in MIBC patients

*ARID1A* is the defining component of the BAF ATP-dependent SWI/SNF nucleosome remodelling complex. In some types of cancer, e.g. ovarian cancer, both deletions and heterozygous truncating mutations result in BAF-complex destabilization and loss of ARID1A protein expression[22]. When evaluating the TCGA MIBC dataset[4], we demonstrated lower ARID1A protein levels in *ARID1A* mutated tumors (P = 0.03, two-sided Wilcoxon-rank sum test), and in tumors harboring *ARID1A* deletions (P = 0.08, two-sided Wilcoxon-rank sum test) (Figure 1A). In addition, positive correlations were detected between ARID1A protein expression, mRNA expression, and copy number status (Supplemental Figure 1A-D).

**Figure 1.**
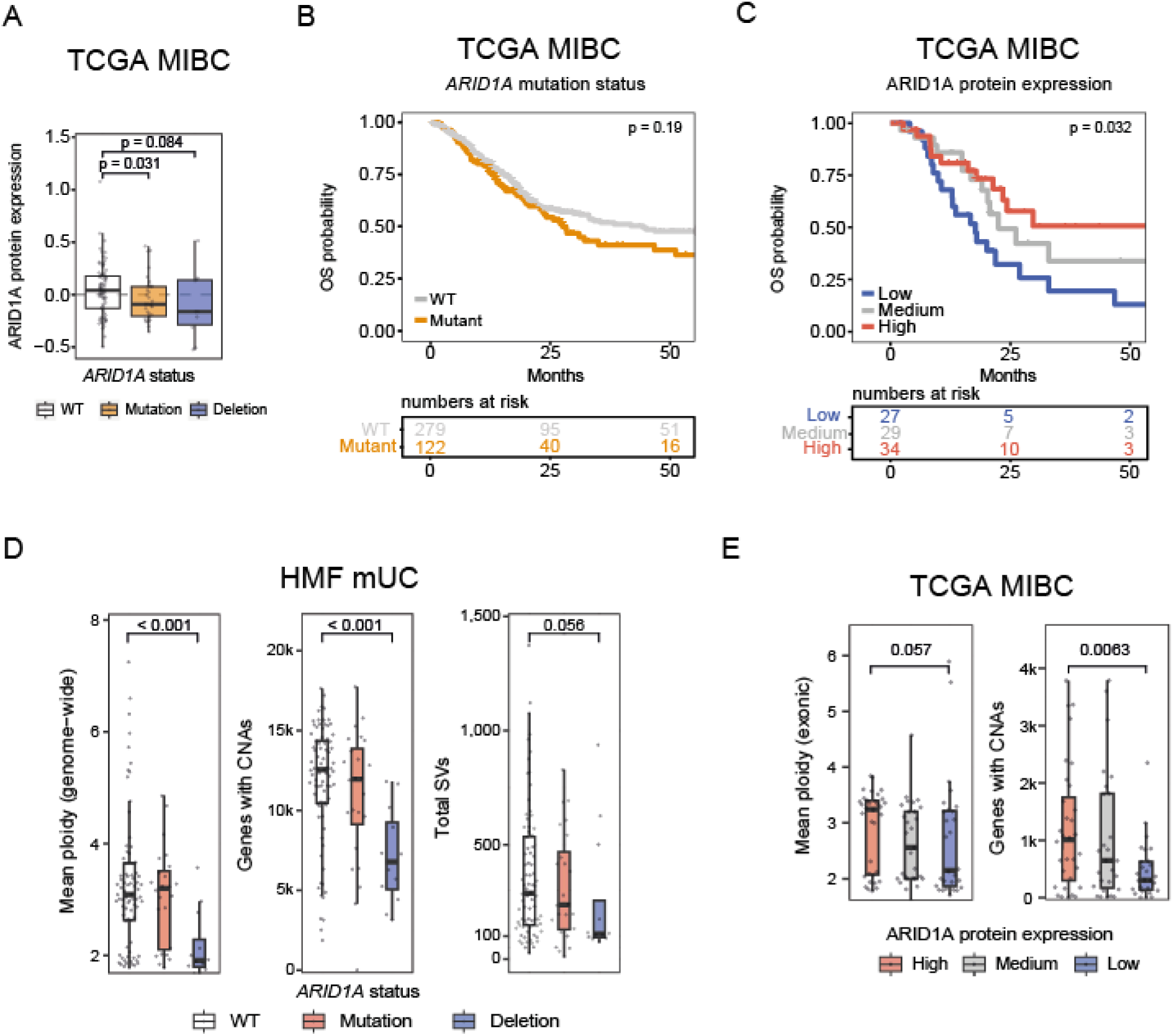
Genetic *ARID1A* aberrations are associated with loss of gene-expression and poor prognosis in MIBC. **A)** Box plots showing ARID1A protein expression levels quantified by reverse phase protein array (RPPA) in *ARID1A* wild type, mutated and deleted tumors*. A two-sided Wilcoxon-rank sum test was applied. Box plots show the median, inter-quartile range (IQR: Q1–Q3) and whiskers (1.5xIQR from Q3 to the largest value within this range or 1.5xIQR from Q1 to the lowest value within this range). WT = *ARID1A* wild type, Mutation = coding mutations (excluding synonymous) and small insertions/deletions, Deletion = *ARID1A* deleted. **B)** Overall survival curves of n = 401 MIBC patients treated with radical cystectomy*. Patients were stratified by *ARID1A* mutation status. WT = *ARID1A* wild type, Mutant includes protein-coding mutations, small insertions/deletions, and *ARID1A* deletions. The log-rank test was applied to survival curves. **C)** Overall survival curves of n = 90 MIBC patients (one patient excluded with no survival record) treated with radical cystectomy*. Patients were stratified by tertiles of ARID1A protein expression levels (low, medium, high) as determined by RPPA and samples with protein-coding mutations were excluded. The log-rank test was applied to survival curves. **D)** Boxplots depicting mean ploidy, number of genes affected by copy-number aberrations (CNAs) and number of structural variants (SV) in metastatic urothelial cancer samples**. WT = *ARID1A* wild type, Mutation = protein coding mutations (excluding synonymous) and small insertions/deletions, Deletion = *ARID1A* deleted. Two-sided Wilcoxon-rank sum test was applied. **E)** Boxplots graphs depicting mean ploidy and number of genes affected by copy-number aberrations (CNAs) in MIBC patients stratified by ARID1A protein expression*. Two-sided Wilcoxon-rank sum test was applied. WT = *ARID1A* wild type, Mutation = protein-coding mutations (excluding synonymous) and small insertions/deletions, Deletion = *ARID1A* deleted. *TCGA **HMF.

We did not observe overall survival (OS) differences when patients were stratified based on *ARID1A* somatic mutation status (p = 0.19, log-rank test) (Figure 1B). However, OS was associated with ARID1A expression at the protein level. Patient stratification into three groups based on ARID1A protein expression (low, medium, high), revealed the shortest OS in patients with low ARID1A protein expression (17.7 months), followed by patients with intermediate expression (22.5 months), while the longest OS was observed in patients with high ARID1A-expressing tumors (59.3 months) (p < 0.05, log-rank test) (Figure 1C), suggesting that attenuated ARID1A protein expression may represent the most clinically relevant indicator of *ARID1A* deficiency. ARID1A protein expression status was associated with worse outcome regardless of treatment, tumor stage, age, or gender (Supplemental Figure 1E). These analyses, highlight the clinical need to develop alternative treatment options for *ARID1A*-deficient patients, who have a poor outcome with current standard of care treatment.

Given *ARID1A*’s association with DNA-repair, we explored whether *ARID1A* mutation status associated with tumor mutational burden. First, we investigated whole-genome DNA sequencing data of an mUC cohort from the Hartwig Medical Foundation (HMF). In this HMF dataset, the number of single-nucleotide variants (P=0.44, two-sided Wilcoxon-rank sum test), indels (P=0.73, two-sided Wilcoxon-rank sum test), or multi-nucleotide variants (P=0.67, two-sided Wilcoxon-rank sum test) per megabase did not differ with respect to *ARID1A* mutation status (Supplemental Figure 1F). Strikingly, when evaluating gross chromosomal aberrations, we found *ARID1A* deletions to be associated with lower overall ploidy (P<0.001, two-sided Wilcoxon-rank sum test), fewer genes affected by copy-number alterations (P<0.001, two-sided Wilcoxon-rank sum test), and fewer structural variants (P=0.056, two-sided Wilcoxon-rank sum test), when compared to *ARID1A* wild type status (Figure 1D). Although whole-genome sequencing data from the HMF cohort offers higher resolution to detect aneuploidy at the gene level than microarrays[23], we observed a similar trend in the TCGA cohort. In the TCGA dataset, patient tumors with low ARID1A protein expression had lower overall ploidy (P=0.057, two-sided Wilcoxon-rank sum test), and fewer genes affected by copy-number alterations (P=0.0063, two-sided Wilcoxon-rank sum test), compared to tumors with ARID1A expression (Figure 1E). Although counterintuitive, these results align with a previous study which demonstrated that *ARID1A* inactivation impairs genome maintenance to such extent that *ARID1A*-deficient cancer cells are vulnerable to DNA double-strand breaks during mitosis, resulting in selective elimination of clones that accumulate too much DNA-damage while dividing [24]. However, the molecular mechanisms underlying the observed prognostic and mutational differences associated with ARID1A status in BC have yet to be elucidated with functional studies.

### Biobank of patient-derived BC tumoroid and normal urothelial organoid models that phenotypically and genetically resemble the tissue of origin

To investigate the impact of ARID1A status on BC in a clinically relevant BC model system, we generated *ex vivo* cultures from BC and macroscopic normal urothelium acquired from BC patients undergoing transurethral resection of bladder tumor (TURBT) or radical cystectomy (Supplemental Figure 2A-C). Patient-derived tumoroid (PDT) and normal organoid (NO) lines were initiated from a variety of BC patients, ranging from low-grade non-invasive NMIBC to high-grade MIBC (Supplemental Figure 2B,D, Supplemental Table 1). Approximately 80% of initiated cultures could be expanded and were successfully cryopreserved (Supplemental Figure 2E-F).

Genome-wide copy-number aberrations (CNA), and hotspot mutation analysis (SNaPshot[25–27]) of the telomerase reverse transcriptase (*TERT*) promoter region and protein-coding regions of Phosphatidylinositol-4,5-Bisphosphate 3-Kinase Catalytic Subunit Alpha (*PIK3CA*) and Fibroblast Growth Factor Receptor 3 (*FGFR3)* confirmed that BC tumoroids and non-malignant bladder organoids genetically resembled their corresponding parental tissues *ex vivo* (Figure 2A-C, Supplemental Figure 3A-B). Additionally, *ex vivo* cultures resembled patient tumor and normal tissue urothelial differentiation marker expression and morphological growth patterns (Figure 2D & Supplemental Figure 3C-D).

**Figure 2.**
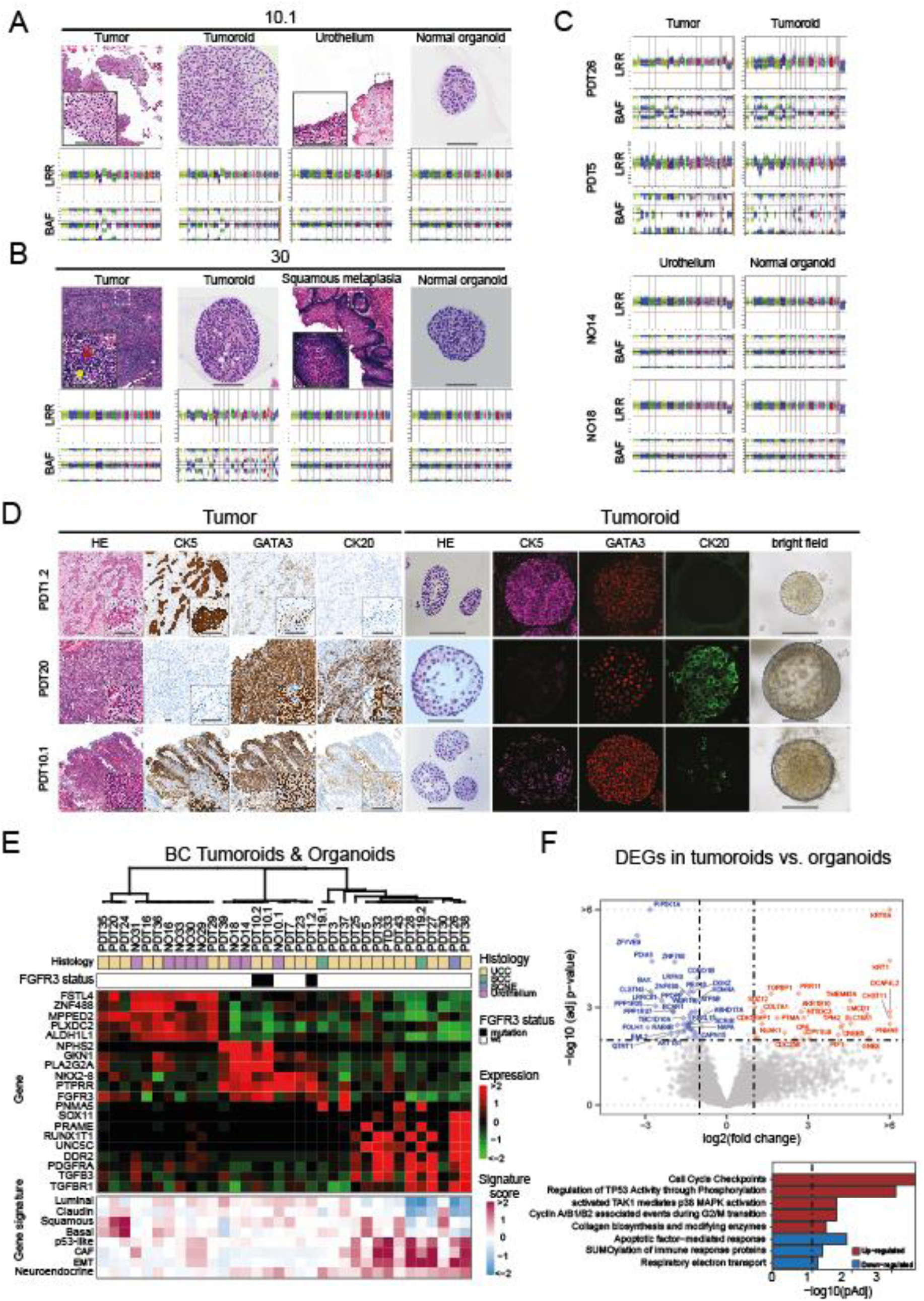
Genetic and phenotypic characterization of a BC tumoroid and normal urothelial organoid biobank. **A)** Top: H&E stainings of patient #10 tissues and corresponding PDTs and NOs (scale bar = 50 μm). Bottom: Scatterplots illustrating genome wide copy number alterations depicted by Log R ratios (LRR) and B-allele frequency (BAF) from #10 tumor (first left), tumoroids (second left), normal urothelium (second right), and normal organoids (first right) **B)** Top: H&E stainings patient #30 tissues and corresponding PDTs and NOs. Red arrow indicates tumor cells, yellow arrow indicates tumor infiltrating lymphocytes (scale bar = 50 μm) Botom: Scatterplots illustrating genome wide copy number alterations from patient #30 tumor (first left), tumoroids (second left), squamous metaplasia (second right), and normal organoids (first right). Note that copy-number alterations (CNAs) are masked in this lymphoepithelioma-like tumor due to high lymphocyte infiltration, but become apparent as CNA resolution increases in the tumoroids. **C)** Scatterplots illustrating genome wide copy number alterations depicted by Log R ratios (LRR) and B-allele frequency (BAF) from patient tumor and corresponding tumoroids (top) or urothelium and corresponding normal organoids (bottom). **D)** Histological evaluation of primary tumors and corresponding tumoroids. Expression of urothelial differentiation markers was investigated by IHC (tumors) and IF (tumoroids) as indicated. Representative examples of basal (PDT1.2; CK+, CK20-), luminal (PDT20; CK5-, CK20+) and mixed basal/luminal (PDT10.1 CK5+, CK2-+) tumor types are shown. (scale bar = 50 μm). **E)** Unsupervised clustering of transcriptomic profiles from patient-derived tumoroids (PDTs) and normal organoids (NOs) identified three clusters (Euclidean distance, Ward method), which are displayed in the dendrogram. Differential gene expression analysis was applied for each group and expression of the top five genes with the highest log2 fold change and adjusted p < 0.001 were shown in the heatmap, together with FGFR3, TGFB3, TGFBR1, PDGFRA and DDR2. Additionally, histological subtypes, FGFR3 mutation status and gene expression signatures of patient-derived tumoroid and normal organoid lines are displayed. Histology represents urothelial cell carcinoma (UCC), small cell neuroendocrine-like (SCNE), squamous cell carcinoma (SCC) and normal urothelium. Tumors with somatic mutations in FGFR3 identified by SNaPshot mutation analysis are indicated in black. Gene signature scores represent the average expression of genes associated with each signature. **F)** Volcano plot of differentially expressed genes obtained from RNA sequencing analysis of tumoroid compared to bladder organoids. Genes that were differentially regulated in tumoroids with adjusted p < 0.01 and absolute log2 fold change >1 are labeled in red (upregulated) and blue (downregulated), (top). Bar diagrams specify the pathways of differentially expressed genes (adjusted p < 0.05, absolute log2 fold change > 0.5) according to the hypergeometric distribution calculated with ReactomePA (adjusted p < 0.05, bottom). P values were adjusted with the Benjamini-Hochberg method.

Unsupervised hierarchical clustering of 3’ mRNA-seq data of BC tumoroid and normal bladder organoid cultures (n = 34) identified three expression subtypes, highlighted by the five most differentially expressed genes (Figure 2E, Supplemental Figure 4A-B). Cluster one contained tumoroids and normal organoids showing a basal/squamous gene expression signature, whereas cluster two seemed enriched for tumoroids and organoids with a luminal expression signature and presence of activating *FGFR3* mutations. The third cluster exclusively consisted of tumoroids and was characterized by low expression of luminal genes and an epithelial-to-mesenchymal transition (EMT)-associated expression signature.

Differential gene-expression analysis of BC tumoroids and normal organoid cultures identified pathways that reflect several hallmarks of cancer and are known to be dysregulated in BC[28]; these include upregulation of pathways involved in MAPK activation, cyclin-associated events during G2/M transition and cell cycle checkpoints (sustaining proliferative signaling), P53 regulation (evading growth suppressors), and collagen modifying enzymes (activating invasion and metastasis), as well as downregulation of pathways promoting apoptosis (resisting cell death) and respiratory electron transfer (deregulating cellular energetics) (Figure 2F). From the present analysis we conclude that patient-derived BC tumoroids and normal urothelial organoids faithfully represent the tissue of origin.

### Loss of *ARID1A* in normal bladder organoids induces upregulation of DNA repair and cell cycle-associated genes

Given the need for novel treatment strategies for patients with loss of ARID1A protein expression, we applied our PDO model to identify potential therapeutic targets that could be exploited in *ARID1A*-deficient BC. Since ARID1A loss has been shown to induce oncogene expression in murine urothelium[29], we investigated which genes are upregulated in response to ARID1A loss in human urothelial cells. Loss of ARID1A expression was modeled by applying short hairpin RNA (shRNA) – mediated ARID1A knock-down in four normal organoid lines (Figure 3A). Efficient *ARID1A* knock-down was confirmed by RT-qPCR (Figure 3B). Thereafter, 3’ mRNA-seq and differential gene-expression analysis identified 353 dysregulated transcripts in ARID1A depleted (N=4) compared to matched parental (N=2), or shCTRL (N=2) control organoids (Figure 3C). A total of 247 genes were significantly upregulated, while 106 genes were significantly downregulated. To explore targetable therapeutic approaches, we concentrated on the upregulated genes, as these present opportunities for inhibition with potential drug treatments. Pathway enrichment analysis pointed to significant upregulation of DNA damage repair (DDR) genes such as *ATM*, *RAD51*, *PALB2*, *BRCA1*, *BARD1*, *FANCA*, *FANCI*, and *FANCG*, and genes involved in cell cycle checkpoint and G1 to G1/S phase transition including *CHEK1*, *CHEK2*, *PRC1*, *BIRC5*, *CDCA8* and *AURKB* (Figure 3C-D). Next, we performed a motif enrichment analysis to identify transcription factors underlying the observed transcriptomic changes associated with ARID1A loss. Homer motif analysis predicted only one de novo motif enriched in the transcription start sites (TSS) of upregulated genes with confidence. This motif is similar to early region 2 factor (E2F) family member binding sites, with the E2F8 binding motif showing the highest resemblance (Figure 3E, Supplemental Figure 5A). The E2F transcription factor family consists of transcriptional activators (E2F1-3) and repressors (E2F4-8) that orchestrate gene-expression in cell cycle regulation and DNA stress response [30–34]. Interestingly, the consensus binding site for the transcriptional repressor E2F8 was found enriched in TSSs of 32 dysregulated genes, of which 25 were upregulated in the ARID1A depleted organoids, pointing to reduction of E2F8 controlled repression upon ARID1A loss (Supplemental Figure 5B). Cross-referencing to publicly available ARID1A CUT&Tag[29] and E2F8 ChIP-seq data[35] demonstrated enrichment of ARID1A and E2F8 binding at the TSSs of genes upregulated by ARID1A knock-down, suggesting a direct link between loss of ARID1A expression and de-repression of E2F8-regulated genes (Figure 3F).

**Figure 3.**
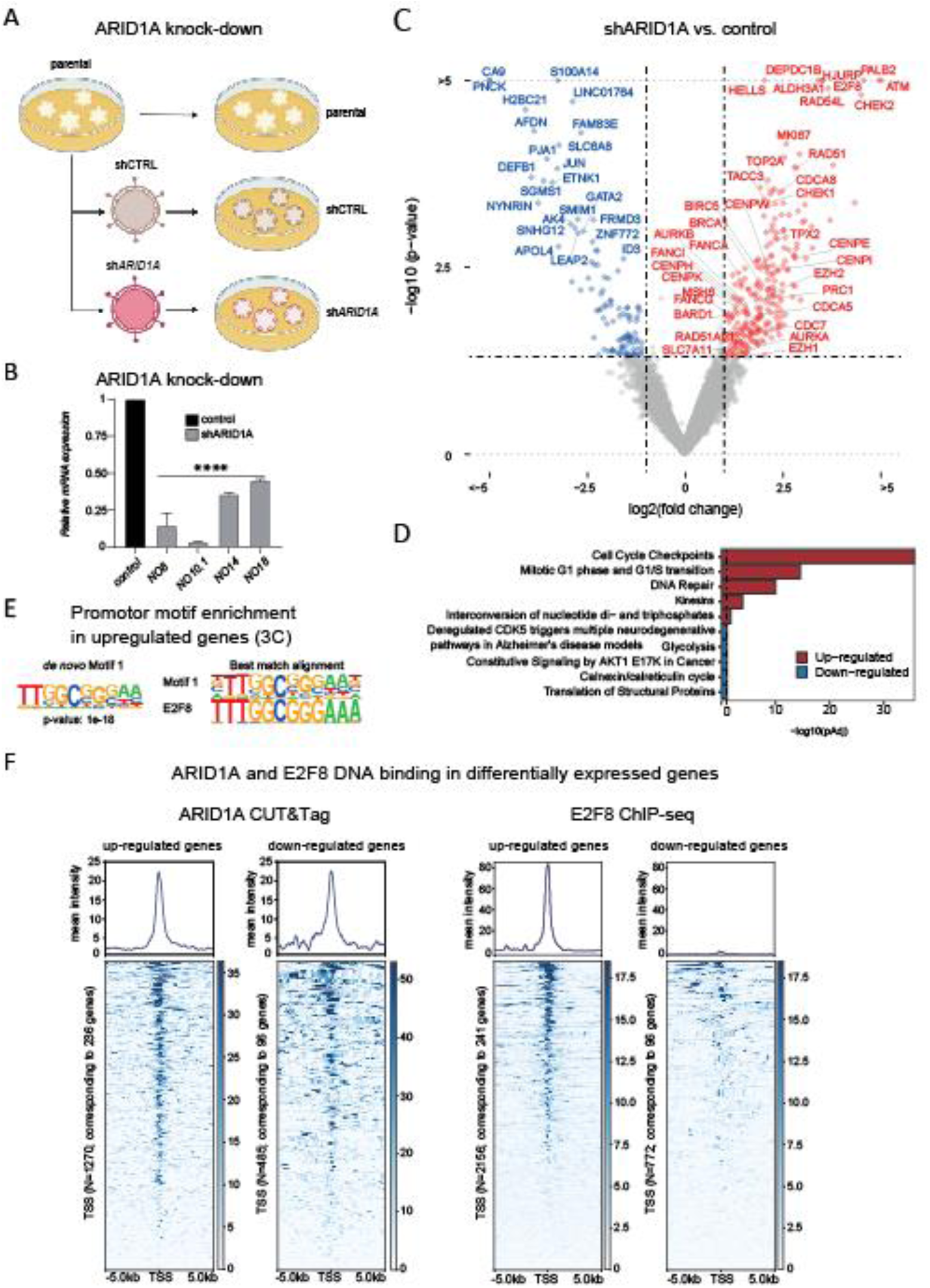
Loss of ARID1A expression in normal bladder organoids induces upregulation of DNA-repair pathways and cell cycle-associated genes. **A)** Schematic representation of the experimental procedure for ARID1A knock-down in urothelial organoids. Following transduction with a lentiviral vector expressing a scrambled shRNA or shRNA targeting *ARID1A* (sh*ARID1A*), organoids were selected with puromycin for 5 days to obtain bladder organoid lines depleted of ARID1A. Four *ARID1A* knock-down (sh*ARID1A*) and corresponding control (shCTRL; NO8 & NO10.1) or untransduced parental (NO14 & NO18) organoid lines were established according to the schematic. **(B)** Levels of *ARID1A* mRNA expression were evaluated by reverse transcription polymerase chain reaction (RT-PCR) in control (shCTRL; NO8 & NO10.1) or untransduced parental (NO14 & NO18), and the transduced (sh*ARID1A*) bladder organoid lines. Expression of *ARID1A* was calculated according to the 2ΔCt method comparing knock-down vs matched controls, using the housekeeping gene Cyclophillin A as reference. One-way ANOVA comparing knock-down vs. control. *** P<0.0005 **C)** Volcano plot of differentially expressed genes obtained from RNA sequencing analysis of control versus ARID1A depleted bladder organoids as indicated. The expression levels of 353 genes were differentially regulated by ARID1A depletion (up- and down-regulated genes are depicted in red and blue, respectively). The name of selected genes is highlighted. P-values were not adjusted for multiple testing. **D)** Bar graph depicting pathways of the differentially expressed genes according to the hypergeometric distribution calculated with ReactomePA (adjusted p < 0.05). **E)** Left: Logo depicting the top *de novo* binding motif (Motif 1), enriched in the promoters of genes upregulated upon ARID1A knock-down (upregulated genes from panel 3C), based on Homer analysis. Right: Logos for predicted Motif 1 and its best match, as determined by sequence alignment (E2F8). **F)** Left: histograms depicting mean ARID1A or E2F8 binding intensities around transcription start sites (TSS) of up- and down-regulated genes as indicated (dysregulated genes from panel 3C). ARID1A CUT&Tag data from murine urothelial organoids was repurposed from Jana et al.[29]. Only genes with murine orthologues (236 out of 247 upregulated genes) are shown. E2F8 ChIP-seq data from K562 (myeloid progenitor was repurposed from the ENCODE project[35]).

### Upregulation of chromosomal passenger complex members and cell cycle checkpoint kinases in *ARID1A*-deficient BC

Cell cycle checkpoint-associated genes were the most significantly upregulated gene set in ARID1A-depleted bladder organoids compared to controls (Figure 3D). The same pattern was observed when comparing tumoroids to bladder organoids (Figure 2F), hereby prompting us to focus on cell cycle checkpoint genes. Ensuring clinical relevance, we cross-referenced upregulated genes identified by ARID1A knock-down to genes upregulated in BC patients with *ARID1A* mutated/deleted or low *ARID1A* expressing tumors. First, we investigated which cell cycle checkpoint genes were significantly upregulated in *ARID1A*-deficient MIBC samples from the TCGA cohort. We identified a total of 73 cell cycle checkpoint-associated genes to be significantly upregulated in MIBC tumors with *ARID1A* deletions (1) and low mRNA expression (72) (Supplemental table 2). Comparing this list to the cell cycle checkpoint-genes upregulated in ARID1A-depleted organoids resulted in 16 overlapping candidate genes (Figure 4A). Unsupervised hierarchical clustering using these genes yielded a clear separation between sh*ARID1A* and control organoids (Figure 4A). Additionally, direct binding of ARID1A and E2F8 was observed at the transcription start sites of the majority of these genes (Supplemental Figure 6). To confirm association of ARID1A status with cell-cycle checkpoint candidate expression, we performed unsupervised hierarchical clustering of the 16 cell-cycle-checkpoint candidate genes on RNA-sequencing data from our patient-derived tumoroids. BC tumoroids separated into two major clusters, suggesting separation by ARID1A status in a manner similar to that observed for ARID1A-depleted organoids (Figure 4B). We randomly selected five BC PDTs from each of the two major clusters and examined ARID1A protein expression using IHC (Figure 4C). IHC identified five BC tumoroid lines with ARID1A expression and five lines without, with eight lines matching the ARID1A status suggested by unsupervised clustering. We then explored whether our 16 cell-cycle-checkpoint candidate genes were also upregulated in *ARID1A*-deficient mUC samples, and whether we could identify *ARID1A*-deficient BC patients by assessing ARID1A protein expression by IHC. For this purpose, we selected samples from the HMF mUC cohort[7, 36] with formalin-fixed paraffin-embedded (FFPE) blocks available for further analysis (N=8). We performed IHC for ARID1A, and summarized *ARID1A* mutation status, ARID1A protein expression and cell-cycle-checkpoint candidate gene mRNA expression into a single heatmap (Figure 4D-E). Unsupervised hierarchical clustering separated the samples into two clearly distinct clusters, one with cell-cycle-checkpoint candidate gene upregulation, and one without. The cluster with upregulation of the 16 cell-cycle-checkpoint candidate genes was enriched for patients with low ARID1A protein expression (Q-score <100, 3 out of 4), *ARID1A* deep deletions (1 out of 4), or *ARID1A* mutations (1 out of 4). Thus, *ARID1A*-deficiency is associated with upregulation of the 16 cell-cycle-checkpoint candidate genes in bladder organoids, BC tumoroids, as well as MIBC and mUC patient tumors.

**Figure 4.**
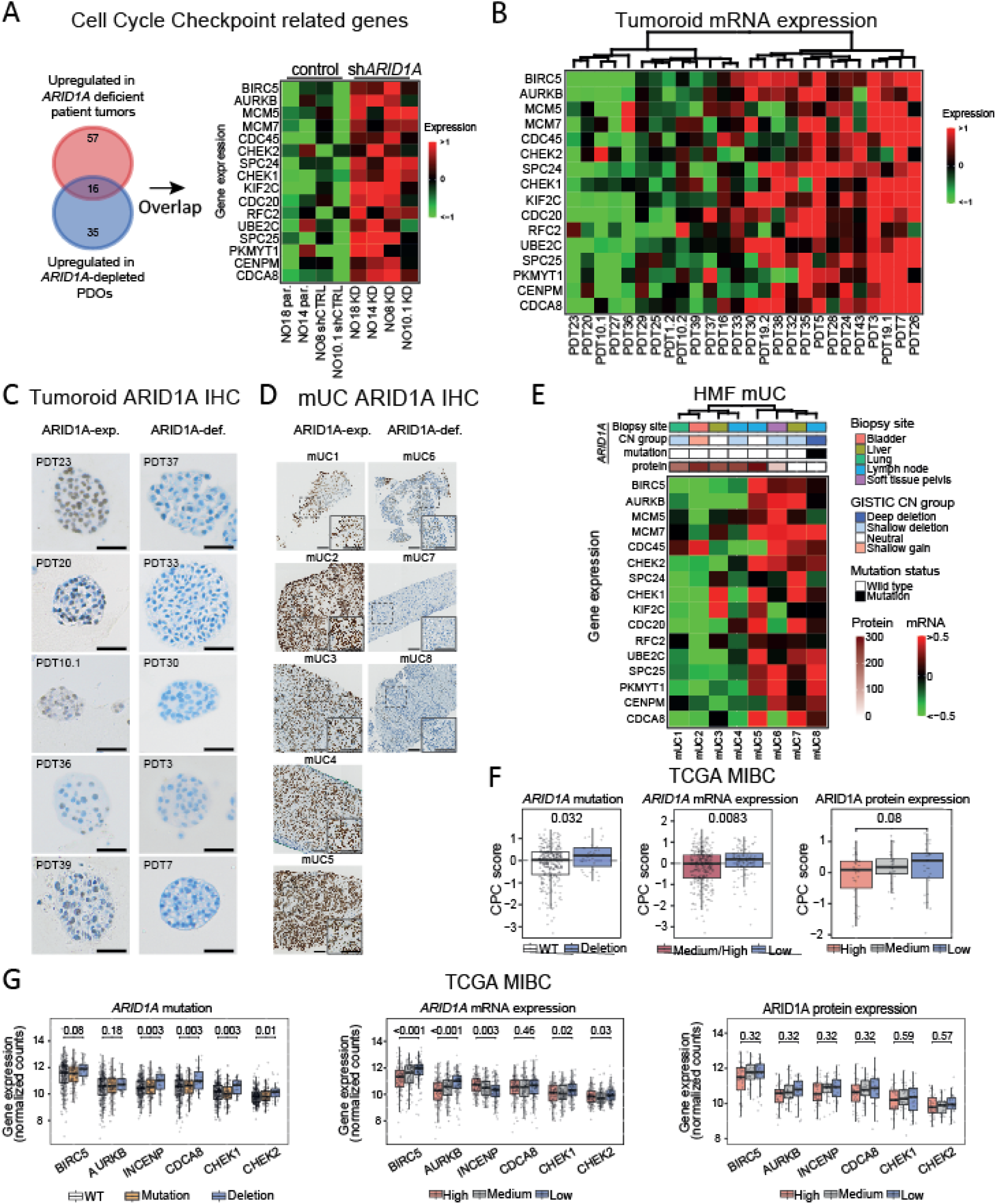
Chromosomal passenger complex members and cell cycle checkpoints are upregulated in *ARID1A*-deficient tumors. **A)** Venn-diagram comparing cell cycle checkpoint genes significantly upregulated in MIBC patient tumors with *ARID1A* deletions or low mRNA* (top) to genes significantly upregulated upon ARID1A depletion in normal urothelial organoids (bottom). Overlapping genes were subsequently used for unsupervised hierarchical clustering (Euclidean distance, Ward method) of ARID1A knock-down (KD), parental (par.) or control (shCTRL) organoid lines. **B)** Heatmap depicting unsupervised hierarchical clustering of BC patient-derived tumoroids (PDTs) by normalized expression of N = 16 cell cycle checkpoint genes associated with *ARID1A* deficiency. **C)** Representative images of ARID1A immunohistochemistry in PDTs and matched patient tumors. (scale bar = 50 μm). **D)** Representative examples of ARID1A-stained metastatic lesions. (scale bar = 100 μm). **E)** Unsupervised hierarchical clustering (Euclidean distance, Ward method) of N = 8 metastatic BC samples summarizing biopsy sites**, *ARID1A* copy-number status**, *ARID1A* mutations status**, ARID1A protein expression investigated by IHC (Q-score), and = 16 cell cycle checkpoint genes associated with loss of *ARID1A*. Data was repurposed from TCGA (*) or HMF (**) datasets. **F)** Boxplots comparing chromosomal passenger complex (CPC) score integrating *BIRC5*, *AURKB*, *CDCA8*, and *INCENP* mRNA expression in MIBC patient tumors with *ARID1A* wild type vs. deleted tumors* (left), medium/high vs. low *ARID1A* mRNA expression* (middle), and high vs. medium vs. low ARID1A protein expression* (right). Two-sided Wilcoxon-rank sum test was applied to compare differences between the groups. **G)** Boxplots comparing mRNA expression of *BIRC5, AURKB, INCENP, CDCA8*, *CHEK1* and *CHEK2* in MIBC patient tumors stratified by *ARID1A* mutation status* (top), mRNA expression levels* (middle) or protein expression levels* (bottom). Two-sided Wilcoxon-rank sum test was applied and p-values were corrected with the Benjamini–Hochberg method. Box plots show the median, inter-quartile range (IQR: Q1–Q3) and whiskers (1.5xIQR from Q3 to the largest value within this range or 1.5xIQR from Q1 to the lowest value within this range). *HMF, ** TCGA.

Three of the identified candidate genes, *BIRC5, CDCA8*, and *AURKB*, form, together with *INCENP*, the chromosomal passenger complex (CPC). The CPC has been described as a master regulator of mitosis, functioning in chromosome–microtubule attachment, activation of the spindle assembly checkpoint, and cytokinesis[37]. The combined mRNA expression of these CPC members was found to be higher in *ARID1A*-deleted and low *ARID1A*-mRNA expressing MIBC tumors, and in patient tumors with low ARID1A protein expression (Figure 4F). Candidate genes *CHEK1* and *CHEK2* are two functionally—but not structurally— related serine/threonine kinases that are activated in response to DNA damage during mitosis. Upon activation, *CHEK1* and *CHEK2* delay cell cycle progression and facilitate DNA repair until damage has been restored, making *CHEK1* and *CHEK2* interesting candidate genes for further investigation[38]. In addition to CPC members, also *CHEK1* and *CHEK2* were significantly upregulated in *ARID1A*-deleted or low *ARID1A*-mRNA expressing MIBC tumors (Figure 4G). From this analysis, we conclude that ARID1A-deficient BC is associated with the upregulation of CPC members (BIRC5, AURKB, INCENP, and CDCA8) as well as checkpoint kinases (CHEK1 and CHEK2).

### Pharmacological *BIRC5* and *CHEK1* inhibition is associated with increased DNA damage signaling in ARID1A-deficient BC

Next, we investigated the effects of pharmacological inhibition of upregulated candidate genes on cell cycle distribution and DNA-damage signaling in context of ARID1A-deficient BC. Clinically advanced inhibitor drugs are available for *BIRC5* (Baculoviral IAP Repeat Containing 5) and *CHEK1* (Checkpoint Kinase 1). *BIRC5* transcription is inhibited by YM155 (sepantronium bromide), leading to decreased expression of the *BIRC5*-encoded protein survivin[39–42]. CHEK1 kinase activity is inhibited by prexasertib, a drug which was recently given FDA fast-track designation for treatment of ovarian and endometrial cancer patients[43]. To gain mechanistic insight into cell cycle distribution and DNA-damage signaling, we used a flow cytometry approach to explore cell cycle distribution and the presence of DNA-damage, both top enriched pathways revealed by our ARID1A-depletion experiments (Supplemental Figure 7A). Cell cycle distribution was similar between ARID1A-expressing (N=3) and ARID1A*-*deficient (N=3) tumoroids, and treatment with prexasertib (*CHEK1*) or YM155 (*BIRC5*) did not induce significant changes between the two groups (Figure 5A, Supplemental Figure 7B). DNA-damage signaling (yH2Ax) was similar between untreated ARID1A-expressing (N=3) and ARID1A-deficient (N=3) BC tumoroids; however, prexasertib or YM155 treatment significantly increased DNA-damage signaling in ARID1A-deficient as compared to ARID1A-expressing tumoroids (Figure 5B-C, Supplemental Figure 7B-C).

**Figure 5.**
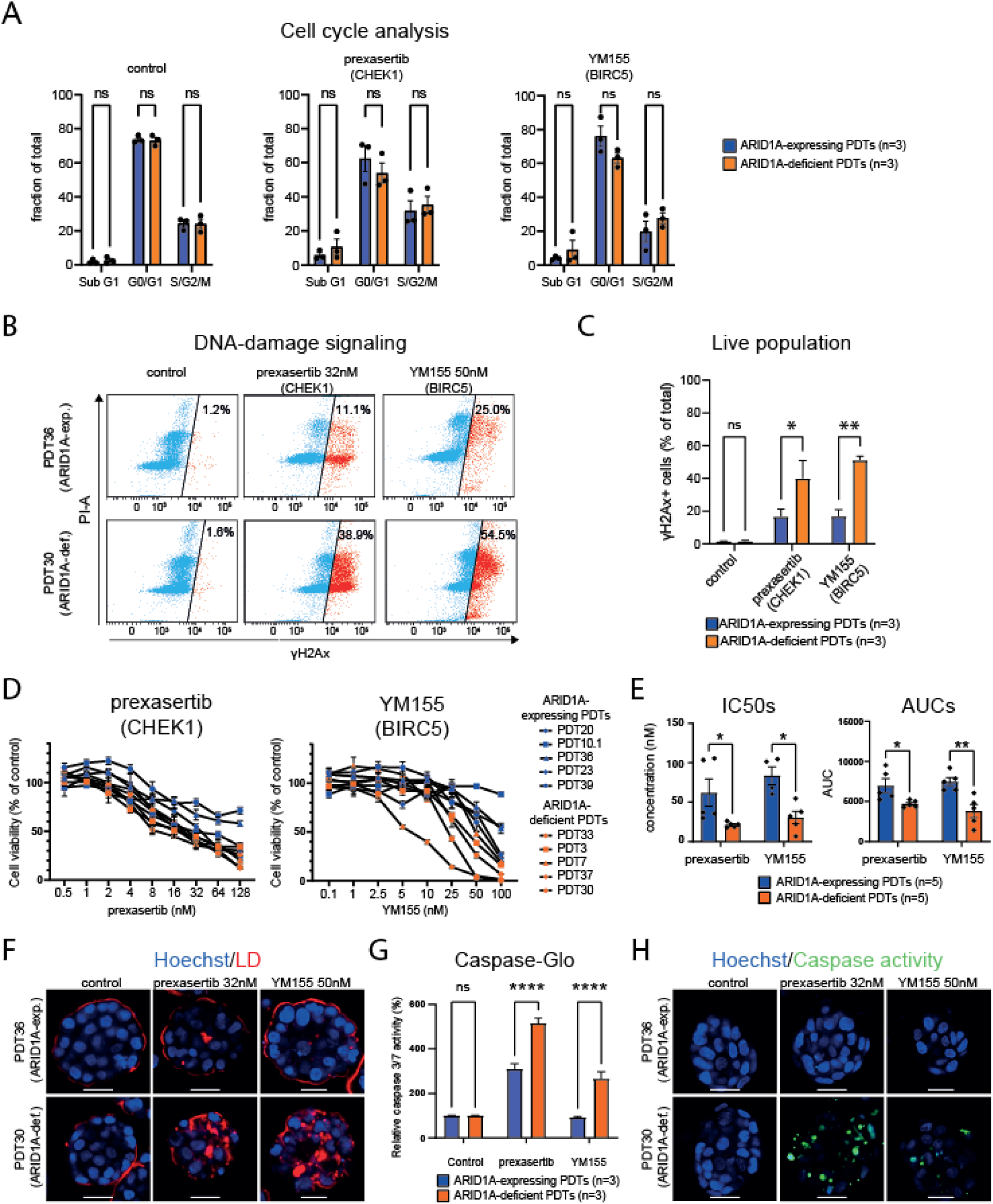
Pharmacological BIRC5 or CHEK1 inhibition selectively targets ARID1A-deficient BC. **A)** Bar graphs summarizing cell cycle distribution of ARID1A-expressing (PDT20, PDT36, PDT39) and ARID1A-deficient (PDT3, PDT30, and PDT37) tumoroids at baseline (untreated control, left), and after treatment with 32nM prexasertib (middle) or 50nM YM155 (right). Data are represented as mean ± SEM. **B)** Flow cytometry of ARID1A-expressing (PDT36) and ARID1A-deficient (PDT30) tumoroids treated with 32nM prexasertib or 50nM YM155 for four days. Gated on the single, live population followed by analysis of propidium iodide (cell cycle) and phosphorylated H2Ax (yH2Ax) to identify the yH2Ax+ fraction (red). **C)** Bar graphs depicting percentage of yH2Ax cells in ARID1A-expressing tumoroids (PDT20, PDT36, PDT39) and ARID1A-deficient tumoroids (PDT3, PDT30, PDT37), treated with 32nM prexasertib or 50nM YM155 for two days. (mean ± SEM, two-way ANOVA, *p<0.05, **p<0.005). **D)** Dose-response curves for ARID1A-expressing (blue) and ARID1A-deficient (orange) PDTs treated with *CHEK1*-inhibitor prexasertib (left), and BIRC5-inhibitor YM155 (right). Cell viability was measured through CellTiter-Glo 3D. (technical triplicates of two independent experiments; mean ± SEM). **E)** Left: IC50 values estimated from non-linear fit of dose-response depicted in D. YM155 IC50 concentration could not be determined for PDT10.1 due to the line’s high resistance. Right: Area under the curve (AUC) of dose-response depicted in D. two-way ANOVA, *p<0.05, ** p<0.005. **F)** Fluorescence staining of ARID1A-expressing (PDT36) and ARID1A-deficient (PDT30) PDTs, treated with 32nM prexasertib or 50nM YM155 for two days. Tumoroids were stained with fixable life/dead staining (LD), nuclei were counter stained with Hoechst. (Scale bar = 25 μm). **G)** Bar graph depicting relative caspase 3/7 activity measured by Caspase-Glo in ARID1A-expressing (PDT20, PDT36, PDT39) and ARID1A-deficient (PDT3, PDT30, PDT37) PDTs following treatment with 32nM prexasertib or 50nM YM155 for two days. (triplicates of two independent experiments; mean ± SEM), two-way ANOVA, ***p<0.0005, ****p<0.00005.) **H)** Fluorescence staining of caspase activity reporter on ARID1A-expressing (PDT36) and ARID1A-deficient (PDT30) PDTs treated with 32nM prexasertib or 50nM YM155 for two days, nuclei were counter stained with Hoechst. (Scale bar = 25 μm).

### Selective susceptibility of ARID1A-deficient BC tumoroids to pharmacological inhibition of *BIRC5* or CHEK1

To investigate if CHEK1 and BIRC5 inhibition can selectively kill ARID1A-deficient cancer cells, we treated ARID1A-expressing (N=5) and ARID1A-deficient (N=5) bladder cancer (BC) tumoroids with a concentration range of inhibitors targeting BIRC5 and CHEK1, followed by cell viability and apoptosis assessments. Both YM155 and prexasertib treatment resulted in patient-specific *ex vivo* drug responses as quantified by CellTiter-Glo 3D (Figure 5D). For both drugs, ARID1A-deficient BC tumoroids (N=5) showed significantly lower IC50 values and area under the curve (AUC) compared to ARID1A-expressing (N=5) BC tumoroids (Figure 5E). Furthermore, ARID1A-deficient BC tumoroids (N=1) treated with YM155 or prexasertib exhibited more cell death than ARID1A-expressing tumoroids (N=1) (Figure 5F, Supplemental Figure 8). Additionally, ARID1A-deficient BC tumoroids displayed significantly increased apoptosis following treatment with prexasertib and YM155, compared to ARID1A-expressing BC tumoroids as was determined by a Caspase-Glo 3/7 assay (n=3 for each group, Figure 5G) and visualized by a caspase 3/7 activity reporter (n=1 for each group, Figure 5H, Supplemental Figure 9). These findings indicate that ARID1A-deficient BC tumoroids are selectively eliminated by YM155 or prexasertib treatment.

To conclude, our ex vivo experiments and patient tumor analyses demonstrate that ARID1A deficiency is associated with the upregulation of E2F8 target genes, including cell cycle-associated genes *BIRC5* and *CHEK1*. In turn, pharmacological inhibition of BIRC5 and CHEK1 via clinically advanced small molecules selectively eliminates *ARID1A*–deficient BC. Our data suggest a promising therapeutic strategy for targeting molecular effectors upregulated due to *ARID1A* deficiency, thereby enhancing the selectivity in eliminating *ARID1A*-deficient BC.

## Discussion

Somatic *ARID1A* mutations are detected in approximately 6% of all human cancers [44], with increased frequencies in bladder cancer (∼30%)[4], ovarian clear-cell carcinomas (∼45%)[22], endometrial carcinomas (∼30%) [22], and gastric cancer (∼15%)[45]. Novel treatments exploiting specific molecular changes introduced by *ARID1A* deficiency could benefit many cancer patients. Pharmacological strategies that exploit *ARID1A* deficiency include PRIMA-1 treatment to inhibit glutathione synthesis[46], homoharringtonine (HHT) treatment to inhibit protein synthesis [9], and GSK126 treatment to inhibit Enhancer of zeste homolog 2 (*EZH2*), the catalytic subunit of the polycomb repressive complex 2 (PRC2), which is known to functionally antagonize mammalian SWI/SNF complexes [47, 48].

In this study, we utilize bladder organoids to pioneer the first human ARID1A-depleted urothelium model, which RNA-sequencing analysis demonstrated to recapitulate *ARID1A*-mutated human BC; we then used this model system to identify novel therapeutic targets for *ARID1A*-deficient BC patients. Patient-derived organoids and tumoroids have demonstrated their invaluable utility as models for studying tumor biology[17, 49]. They faithfully mirror the *in vivo* characteristics of their native tissues, and, when combined with modern experimental techniques, offer experimental versatility and a robust platform for investigating the molecular underpinnings of bladder cancer[19, 20, 50–52]. Moreover, PDTs have been shown to predict patient treatment response, as shown for colorectal, ovarian, and pancreatic cancer[15, 53–55]. Genetic transformation of human organoids has provided significant mechanistic understanding of oncogenic loci identified through extensive genome sequencing investigations of human malignancies[12, 56–58]. Using this ARID1A-depleted urothelium organoid model we identified significant upregulation of DNA repair genes, genes involved in cell-cycle-checkpoint and G1 to G1/S phase transition. Interestingly, the consensus binding site for the transcription repressor E2F8 was significantly enriched in the TSSs of the up-, but not in down-regulated genes. This indicates that upregulation of a subset of genes upon ARID1A depletion is (partly) mediated through modulation of E2F8 binding and suggests a model in which ARID1A facilitates binding of E2F8 to its consensus sites to allow transcriptional repression of its target genes (Supplemental Figure 10). This is supported by data from previous studies reporting direct binding of E2F factors by ARID1A[59], ARID1A-dependent repression of E2F-responsive genes[60–62], and E2F8 binding at ARID1A target genes[9, 33, 64]. Many E2F target genes exhibit an oscillatory expression pattern throughout the cell cycle, characterized by low expression during the M/G1 phase, an increase in expression during the G1/S transition, and a decrease during the G2/M phase[63, 64]. E2F target gene expression is orchestrated by transcriptional activators (E2F1-3) and repressors (E2F4-8) competing for similar binding motifs [30–34]. E2F8 target gene repression has been reported critical to induce S-phase arrest in response to DNA damage, allowing DNA repair and maintenance of genetic stability [64, 65]. De-repression of E2F8 targets consequent to *ARID1A*-deficiency could thus lead to an inability to induce s-phase arrest, causing the elimination of tumor cells with replication stress. In turn, the elimination of tumor cells accumulating DNA-damage during cell division aligns with the low number of structural variants and copy number aberrations we observed for *ARID1A* deficient patient tumors (Figure 1, Supplemental Figure 1), and susceptibility of *ARID1A*-deficient tumoroids towards cell cycle checkpoint inhibition (Figure 5).

Similar to the ARID1A-depleted urothelium organoid model, patient-derived tumoroids lacking ARID1A expression and *ARID1A*-deficient MIBC patient tumors also exhibited increased expression of cell cycle checkpoint and DNA repair genes. Prior studies in various malignancies, including BC, have associated *ARID1A* deficiency with impaired DNA double-strand break repair [29, 66, 67] and cell cycle defects [24, 68, 69]. We showed that cell cycle checkpoint inhibition with prexasertib (targeting CHEK1 kinase activity), and YM155 (targeting *BIRC5* transcription) activated apoptotic pathways and cell death in ARID1A*-* deficient, but not ARID1A-expressing tumoroids. Similar observations have been made recently in a study by Lo et al., identifying *BIRC5* as a therapeutic target in *ARID1A*-deficient gastric cancer tumoroids[12]. Both prexasertib and YM155 are currently under clinical investigation. Prexasertib monotherapy demonstrated a mild toxicity profile and durable responses in patients with platinum-relapsed ovarian cancer[70], which, like BC, is characterized by frequent somatic *ARID1A* mutations. Notably, prexasertib was granted FDA fast-track designation for treatment of ovarian cancer patients[43]. Meanwhile, YM155 is investigated in a phase II study enrolling B cell lymphoma patients (NCT05263583). Prior phase I/II studies already concluded that YM155 is well tolerated; however, it lacked clinical efficacy in unselected prostate cancer and B cell non-hodgkin lymphoma patients (29-30). Our *ex vivo* response data, however, suggests that YM155 and prexasertib may be more efficacious in patients with *ARID1A-*deficient tumors. Our demonstration that it is feasible to stratify patients by ARID1A IHC, along with favourable toxicity profiles of YM155 and prexasertib, raises the prospect of rapid translation into clinical trials.

In conclusion, our study of ARID1A-depleted human bladder organoids and ARID1A-deficient BC tumoroids enabled the identification of cellular processes disrupted in the context of *ARID1A* deficiency, highlighting the power of this patient-representative platform in disease modelling. Our analyses provide mechanistic insights into secondary dependencies of ARID1A-deficient BC, and our *ex vivo* validation of top therapeutic candidates employing clinically advanced pharmacological inhibitors holds potential for rapid translation into the clinic. Similar strategies employing oncogene-engineered organoids could be extended to encompass a wider range of cancer-associated genes and various types of tumors, potentially yielding valuable clinically relevant insights into oncogenic transformation and, in the end, therapeutic strategies.

## METHODS

### Organoid/tumoroid culture

Human bladder tissue was obtained from the Erasmus MC Bladder Cancer Center, Rotterdam, the Netherlands, the Amphia Ziekenhuis, Breda, the Netherlands, and the HagaZiekenhuis, The Hague, the Netherlands. Bladder organoids and BC tumoroids from biopsies obtained through TURBT or cystectomy were isolated and cultured using methods developed by Mullenders et al. [20] with modifications. Briefly, bladder tissues were washed with Advanced DMEM/F12 (Gibco) supplemented with 10mM HEPES (Gibco), 1% GlutaMax (Gibco) and 100 μg/ml primocin (InvivoGen), henceforth Ad+++. Tissue was minced and incubated at 37°C with collagenase 2.5mg/ml in EBSS (Gibco) for 60-90 minutes and isolated cells were passed through 70μM strainer (Falcon), washed with Ad+++ and seeded in 50 µl drops of BME (R&D system) containing 10000-15000 cells in 24 well suspension plates (Greiner). Tumoroids and organoids were cultured in a culture medium containing Ad+++ supplemented with 1 × B-27 (Gibco), 1.25 mM N-acetylcysteine (Sigma), 10 mM nicotinamide, 20μM TGFβ receptor inhibitor A83-01, 100ng/ml recombinant human FGF10 (Peprotech), 25 ng/ml recombinant human FGF7 (Peprotech), 12.5 ng/ml recombinant human FGF2 (Peprotech), 10μM Y27632 Rho Kinase (ROCK) Inhibitor (Sigma) and conditioned media for recombinant Rspondin (2.5% v/v), and Wnt3A (2.5% v/v), henceforth bladder organoid medium (BOM). Cultures were passaged at a 1:3 to 1:6 ratio every 7-14 days. For passaging, BME was first digested with 500μg/ml dispase (Gibco, 17105041) for 1 h at 37°C. Cultures were collected in 15mL tubes, Ad+++ was added to 10mL total volume, and organoids were pelleted by centrifugation at 200 x g. Supernatent was discarded, and tumoroids/organoids were dissociated to single cells using cell dissociation solution-non enzymatic (Sigma, C5914) and mechanical dissociation with a P200 pipette. Dissociated single cells were washed once with 10mL Ad+++, centrifugated at 200 x g, and resuspended in a mixture of culture medium and BME in a 1:2 ratio, and dispersed in new drops. Drops were solidified in the incubator at 37°C for 45 minutes, followed by addition of pre-warmed BOM. Medium was changed every three to four days.

### Production of shRNA lentiviral vectors

Lentiviral constructs containing the desired shRNA sequences (shControl - SHC002 and sh*ARID1A* - TRCN0000059089; targeting sequence: GCCTGATCTATCTGGTTCAAT) were amplified from bacterial glycerol stocks obtained in house from the Erasmus Center for Biomics and part of the MISSION® shRNA library. 5.0 x 10^6^ HEK293T cells were plated in a 10 cm dish and transfected with 12.5 μg of plasmids mix. 4.5μg of pCMVΔR8.9 (envelope) [71], 2 μg of pCMV-VSV-G (packaging) [71] and 6 μg of shRNA vector were mixed in 500 μL serum-free DMEM and combined with 500 μL DMEM containing 125 μL of 10 mM polyethyleneimine (PEI, Sigma). The resulting 1 mL mixture was added to HEK293T cells after 15 min incubation at room temperature. The transfection medium was removed after 12 hours and replaced with a fresh RPMI medium. Virus-containing medium was harvested and replaced with fresh medium at 48, and 72 hours post-transfection. After each harvest, the collected medium was filtered through a cellulose acetate membrane (0.45 μm pore), concentrated by ultra-centrifugation and used directly for shRNA transductions.

### Organoid transduction

Organoids were dissociated to single cell applying the same methods as for passaging. Per condition, 500.000 single cells and 1mL concentrated virus were gently mixed and dispersed into two wells of a 24-well plate. Plates were then centrifuged at 600 x g for 1 hour at 32 °C (spinocculation). The organoid/lentivirus was gently mixed using a P1000 pipette, to detach any adherend cells. Plates were incubated for 5 hours at 37°C, after which single cells were collected in falcon tubes, washed with 10mL ad+++ and centrifuged at 250g for 10 min at 4 °C. Supernatant was removed and transduced cells were seeded in pre-warmed suspension plates. Selection started three days after transduction, using 2 µg/ml puromycin (Invivogen, ant-pr). Puromycin was removed after four days and organoids were cultured for 7-10 days until recovery. Knock-down confirmation was performed with RT-qPCR.

### RNA extraction, cDNA synthesis and Real Time-quantitative PCR (RT-qPCR)

Bladder organoids and tumoroids were harvested with dispase, washed once with 10mL Ad+++ and centrifuged for 200 x g for 5 minutes. Then, 1 mL of cold TRIzol (TRI Reagent®, Sigma-Aldrich, T9424) was added to the organoid pellet and the pellet was thoroughly mixed by intermittent vortexing for 1 minute. Samples placed on ice for 10 minutes, prior to storing at -80 C until further processing. RNA was isolated using standard phenol-chloroform RNA extraction. Briefly, frozen samples were equilibrated to room temperature for 15 minutes. Then, 200 µl Phe-nol:Chloroform:Isoamyl Alcohol 25:24:1 (Sigma-Aldrich, P3803) was added and the samples were thoroughly vortexed and incubated at RT for 2 minutes. Samples were subsequently centrifuged at 12000 x g for 15 min at 4 °C. The aqueous phase was transferred to a fresh Eppendorf tube, 500 µl isopropanol (Biosolve Chimie, 220703) was added and samples were incubated for 15 min at RT. Samples were centrifuged (12000 x g, 4 °C, 10 minutes), supernatant was removed and pellets were washed twice with 1 ml 75% ethanol (Honeywell, 32221), centrifuging at 7500 x g for 5 min at 4 °C. As much supernatant was removed as possible, and RNA pellets were air-dried for 20 min at RT. The RNA was dissolved in 20-40 µl nuclease-free water (Promega, P119E), quantified using NanoDrop® Spectrophotometer ND-1000 (Isogen Life Science) and stored at -80 °C until further usage.

cDNA was synthesized from 500-1000 ng RNA using SuperScript™ II Reverse Transcriptase (200 U/µl, Invitrogen, 100004925) according to manufactorers protocol. cDNA samples were diluted to a final concentration of 2.5 ng/µl using nuclease-free water and stored at -20 °C until further usage.

RT-qPCR was performed using GoTaq® qPCR Master Mix (Promega, A6002), according to manufacturer’s protocols, using CFX96™ Real-Time PCR Detection System (Bio-Rad Laboratories, Singapore). PCR experiments included an initial denaturation at 95 °C for 5 min, 40 amplification cycles starting at 95 °C for 10 sec, followed by 60 °C for 30 sec. Melting curves were assessed by complete annealing and gradual increase in temperature from 65 °C to 95 °C. The data were analyzed using 2-ΔΔCt methods [41], expressed as relative gene expression and normalized to the reference gene Cyclophilin A. Negative controls were used in each reaction plate. The following forward and reverse primers used for RT-qPCR were synthesized by Integrated DNA Technologies: Cyclophilin A (forward: TCATCTGCACTGCCAAGACTG; reverse: CATGCCTTCTTTCACTTTGCC), *ARID1A* (forward: GTCTCAGCAGTCCCAGCAAA; reverse: GATA-GATCAGGCAAGCTGGAGG).

### Organoid RNA-seq

Total RNA was isolated and quantified as described above. Quality was assessed on a Bioanalyzer (Agilent Technologies) using the Agilent RNA 6000 Nano Kit reagents. Library preparation was performed using the 3’ mRNA-seq Library Prep Kit for Ion Torrent (QuantSeq-LEXOGEN, Vienna, Austria). The libraries were quantified and pooled together at a final concentration of 100 pM. The libraries were templated and enriched on an Ion Proton One Touch system and templating was performed using Ion PI Hi-Q OT2 200 Kit (ThermoFisher). The sequencing was performed using Ion PI Hi-Q Sequencing 200 Kit on Ion proton PI V2 chips (ThermoFisher). Fastq files were mapped to the Genome Reference Consortium Human Build 37 (GRCh37), using a two-step alignment process. Firstly, reads were mapped with hisat2 [72], using default parameters. Next, the unmapped reads were mapped with bowtie2 [73] using the --local and --very-sensitive parameters. Counting of the reads on the 3’ UTRs was performed with metaseqR2 [74].

Differential expression analysis between the tumoroids and organoids was performed with metaseqR2 using the DESEQ2[75] algorithm with default settings and exonFilters = NULL. Expression was corrected for samples derived from the same donor. Pathway enrichment analysis was performed with ReactomePA v1.44.0 [76] using differentially expressed genes with adjusted p < 0.05 and log2 fold change > 0.5. Significantly enriched pathways were defined as having adjusted p < 0.05 and only the top 5 up- and down-regulated pathways were displayed in the figures. For unsupervised hierarchical clustering, transcript counts were normalized with DESeq2 applying variance stabilizing transformation on protein-coding transcripts. The normalized counts were subsequently median-centered and the Euclidean distance calculated to perform hierarchical clustering with the Ward method. Gene expression signature scores were calculated as the average median-centered expression of genes associated with each signature [7]. Accordingly, for the ARID1A KD comparisons, we employed the PANDORA algorithm within the metaseqR2 package by integrating the DESEQ, DESEQ2, edgeR, limma, NBPSeq, and NOISeq algorithms. Differentially expressed genes were identified based on a meta p-value threshold of < 0.05 and a log2 fold change > 1.

### The Cancer Genome Atlas (TCGA) bladder cancer cohort

The TCGA data for the bladder cancer cohort is publicly available at https://portal.gdc.cancer.gov./. Somatic mutations detected by Mutect of 412 tumors, GISTIC copy number changes at gene level of 410 tumors, ARID1A protein expression quantified by reverse phase protein array of 127 patients and RNA-seq (HTSeq counts; Affymetrix SNP6 arrays) data available for 410 tumors were analyzed. *ARID1A* was considered deleted when gistic score was <-0.4. Transcript counts were normalized with DESeq2 v.1.32.0 applying variance stabilizing transformation on protein-coding transcripts. Samples were stratified in tertiles according to the ARID1A protein expression into n=42 low, n=43 medium and n=42 high expressed groups. For downstream analysis, samples with protein-coding mutations were excluded, resulting in n=28 low, n=29 medium and n=34 high ARID1A expressed groups. RNA counts were used for differential gene expression analysis between *ARID1A* deleted samples vs the rest, low ARID1A protein-expressed samples vs the rest and low *ARID1A* mRNA-expressed group vs the rest. Differentially expressed genes had adjusted p < 0.05 and absolute log2 fold change > 1. Multivariate cox regression analysis was applied using the survival R package[77].

### HMF metastatic urothelial cancer cohort

WGS and RNA-seq data from metastatic urothelial carcinomas are available through the Hartwig Medical Foundation at https://www.hartwigmedicalfoundation.nl/en/data/data-access-request/, under request number DR-314. Samples that were previously analyzed by Nakauma-Gonzalez et al.[7] were retrieved from DR-314 and re-analyzed with the same bioinformatics pipeline using the human reference genome hg19. *ARID1A* was considered deleted when gistic score was <-0.9. RNA counts were normalized with DESeq2 v.1.32.0 applying variance stabilizing transformation on protein-coding transcripts.

### Histology & Histochemistry

Patient tissue was processed using standard procedures. For tumoroid and organoid processing, tumoroids/organoids were fixed within BME-drops using 4% paraformaldehyde + 0.2% glutaraldehyde (in-house produced) at room temperature for two hours. Fixed BME-drops were then pre-embedded in 4% agarose prior to paraffin embedding. H&E staining were done automatically in the HE600 (Ventana). Alcian Blue staining was done automatically according to the manufactures instructions on the Ventana special stains (#860-002, Ventana).

### Immunohistochemistry

Protein expression of UCC differentiation markers and ARID1A was investigated by automated IHC using the Ventana Benchmark ULTRA (Ventana Medical Systems Inc.). Sequential 4 µm thick (FFPE) sections were stained for markers indicated below (Table 1) using Optiview detection (OV) (#760-700, Ventana) or Ultraview detection (UV) (#760-500, Ventana). In brief, following deparaffinization and heat-induced antigen retrieval with CC1 (#950-500, Ventana), the tissue samples were incubated with antibody of interest for the indicated time (Table 1). Incubation was followed OV, UV detection and hematoxylin II counter stained for 8 minutes followed by a blue coloring reagent for 8 minutes according to the manufactures instructions (Ventana).

**Table 1.** Immunohistochemistry information.

| Antibody | Dilution | Species | Company | Catalog and Clone | Pretreatment in minutes | Ab incubation time at 37°C |
| --- | --- | --- | --- | --- | --- | --- |
| GATA3 | 1,63 µg/ml | Mouse | Cell Marque | L50823 | CC1 32' OV | 48 minutes |
| KRT5 | 0,21 µg/ml | Rabbit | Cell Marque | SP27 | CC1 64' UV | 32 minutes |
| ARID1A | 1/1000 | Rabbit | Abcam | EPR13501 | CC1 32' OV | 32 minutes |
| KRT20 | 0,16 µg/ml | Rabbit | Ventana | SP33 | CC1 32' OV | 24 minutes |
| Synaptophysin | 0,06 µg/ml | Rabbit | Ventana | SP11 | CC1 32'OV | 24 minutes |

### SNaPshot mutation analysis

DNA was isolated using with the QIAmp DNA Mini-Kit (Qiagen) according to the manufacturer’s protocol. Presence of hotspot mutations in the *TERT* promoter sequence chr5:1,295,228C>T, chr5:1,295,248G>A and chr5:1,295,250C>T [GRCh37/hg19]), *FGFR3* (R248Q/E, S249C, G372C, Y375C, A393E, K652E/M) and *PIK3CA* (E542K, E545G/K and H1047R) were assessed on tumor, tumoroid, normal adjacent urothelium and organoid DNA by SNaPshot mutation analysis with the same methods as previously described [25–27]).

### Copy-number aberration analysis in tumoroid/organoid samples

Copy number aberration analysis was performed using single-nucleotide polymorphism (SNP) microarrays (Infinium Global Screening Array (GSA) V3, Illumina) on tumor, tumoroid, normal adjacent urothelium and organoid DNA using standard protocols. SNP data (log-R ratio, B-allele frequency) were visualized to identify potential CNVs via Biodiscovery Nexus CN7.5. (Biodiscovery) and the GenomeStudio genotyping module (Illumina).

### Immunofluorescence

Bladder cancer tumoroids/organoids were cultured and stained in chamber slides (ThermoFisher, 154526PK) prior to fixation with 4% paraformaldehyde for 20 minutes at room temperature. Organoids were washed three times with PBS followed by a 30 minute treatment with 0,1M glycine and 30 minute permeabilization with 0.5% Triton X-100 (Sigma) in PBS, both at room temperature. Organoids were blocked with 0.3% Triton-X-100 (Fluka, 93426), 1% DMSO (Honeywell Riedel-de Haën, 34869), and 0.5% goat serum (Vector Laboratories, S 1000), in PBS at room temperature for 2 hours. Following blocking, organoids were incubated overnight with primary antibodies: KRT5 (Bio Legend; 905901), KRT20 (Dako; M7019), GATA3 (Cell Signaling Technology, D13C9). After three washes in PBS, organoids were stained with appropriate Alexa Fluor dye-conjugated secondary antibodies (Invitrogen), 1:1000 for 1 hour at room temperature. Chambers were removed from the slides and slides were mounted with DAPI (Southern Biotech, 0100-20) and a cover slide. Immunofluorescence images were acquired using a confocal microscope (Leica, Stellaris). Brightness and contrast was adjusted in Image J.

### *Ex vivo* drug response

Bladder tumoroids were cultured for 7-10 days in BME prior to harvesting and dissociation to single cells as previously described. Per condition, 10.000 cells were seeded in 100µl bladder organoid medium containing 15% BME. YM-155 (PBS, Selleckchem, S1130) and prexasertib (DMSO, Selleckchem, LY2606368) were added when mature organoids formed after two-three days, and drug treatment was performed in triplicate. Tumoroids were treated with concentrations as indicated, and treatment lasted four days (96h) prior to viability assessment, whereas a two day (48h) treatment was used for other read-outs. Concentrations used for single-dose treatment (prexasertib 32nM and YM155 50nM) were based on average IC50 values of ARID1A-expressing and ARID1A-deficient tumoroid lines. Following treatment, cell viability was assessed by by cellTiter-Glo 3D (Promega, G9681) and Caspase 3/7 activity by Caspase-Glo (#G8093, Promega). Plates were read on a SpectraMax I3 plate reader. Viability data was normalized using organoid wells treated with vehicle control (0.02% DMSO, 1.2% PBS). Presented data are triplicates from two independent experiments.

For fluorescence analysis of caspase 3/7 activity, treated organoids were harvested with dispase, and washed 2 times with Ad+++ to completely remove BME. Organoids were processed in a 96 well flat, clear-bottom microscopy plate (Revvity, 6005225). Organoids where incubated with caspase 3/7 activity dye (CellEvent™ Caspase-3/7 Detection Reagents, 1:1000 in BOM) for 30 minutes at 37°C, followed by fixation with 4% paraformaldehyde at room temperature for 30 minutes. Organoids were washed once with 200µL PBS, followed by staining with 2 µg/mL Hoechst33342 (Molecular Probes) in 200 µL PBS at 4°C for 12 hours. Plate was imaged with the Opera Phenix Plus High-Content Screening System (Revvity, Waltham, MA, USA). Pre-scanning using 10x bright field images was performed to identify fields of view containing organoids. Fields of view containing organoids were then imaged at 40X magnification for confocal and 20x for bright field, covering the center of each well. 25 Fields of view were imaged per condition. Image analysis was performed with the Harmony software (Revvity). Hoechst was imaged excitation 405nm, emission 435-480nm, caspase reporter activity with excitation 488nm, emission 500-550nm. Briefly, organoids containing at least three nuclei were automatically segmented and mean caspase reporter intensity per organoids was calculated for all conditions. Measurements were obtained for at least 49 organoids per condition and statistical analysis was performed using a mixed-effect analysis due to different samples sizes. For image display, raw images were exported and brightness and contrast was adjusted in imageJ.

For fluorescent Live/Dead staining, treated tumoroids were harvested using dispase and washed once with 500µl PBS. Tumoroids were stained with fixable live/dead staining (Invitrogen™L34975) 1:500 in PBS for 15 minutes PBS, following reconstitution according to manufacturer’s protocol (Invitrogen, L34994). Tumoroids were quenched with 1mL PBS + 5% FCS and fixated with 4% paraformaldehyde at room temperature for 30 minutes. Organoids were washed once with 500µL PBS, followed by staining with 2 µg/mL Hoechst33342 (Molecular Probes) in 200 µL PBS at 4°C for 12 hours. Tumoroids were transferred to a 96 well flat, clear-bottom microscopy plate (Revvity, 6005225), and the plate was imaged with the Opera Phenix Plus High-Content Screening System (Revvity). Pre-scanning using 10x bright field images was performed to identify fields of view containing organoids. Fields of view containing organoids were then imaged at 40X magnification for confocal and 20x for bright field, covering the center of each well. 25 Fields of view were images per condition. Hoechst was imaged excitation 405nm, emission 435-480nm, fixable live/dead staining with excitation 488nm, emission 500-550nm. Image analysis was performed with the Harmony software (Revvity). Software was set to measure mean signal intensity of live/dead dye in the inner 60% of the tumoroids, excluding the outer rim in order to limit detection of non-specific staining on the outside of the organoids. For image display, raw images were exported and brightness and contrast was adjusted in imageJ unless stated otherwise.

### Flowcytometry

For flow cytometry, tumoroids were treated with 32 nM prexasertib, 50 nM YM-155, or vehicle control for 48 hours. Tumoroids were harvested with dispase, and dissociated with non-enzymatic dissociation solution. Single cells were washed twice and stained with fixable live/dead staining (Invitrogen™L34975) 1:1000 in PBS for 15 minutes, followed by one wash with PBS + 5% FCS, and fixation in 100% methanol for 20 minutes. Samples were stored at 4°C for up to a week or stained fresh. Antibody staining was performed on ice for 3 hours with mouse anti-γH2AX (4ug/mL) CAT#) in permeabilization and blocking buffer, followed by two washes in permeabilization and blocking buffer. Subsequent secondary staining was done with anti-mouse Alexa 647 (4ug/mL CAT#) for 90 minutes on ice, followed by two washes with permeabilization and blocking buffer. Cells were subsequently stained with 50 µg/mL propidium iodide (PI) (Sigma) and 0.2 mg/mL RNAse A (10109142001, Sigma-Aldrich) for 30 minutes at room temperature. Stained cells were then FACS-analyzed using a 655 LP and a 695/40 BP filter. Events were gated for single, and live cells based on PI intensity and live/dead staining. Tumoroid-specific gates were set to determine the fraction of γH2AX positive cells and cell cycle distribution. Prexasertib and YM155 treatment distorted cell cycle distribution in such extend that a clear distinction between phases S and G2/M was not possible, requiring the merging of S, G2 and M in further analyses. One-way ANOVA was used to compare γH2AX positive fractions and cell cycle distribution between ARID1A-expressing and ARID1A-deficient tumoroids.

### Statistical Analysis

All data from lab experiments are presented as mean ± standard error of the mean (SEM) and are the result of two independent experiments performed in triplicate, unless stated otherwise. Statistical significance was calculated as indicated in figure legends. GraphPad Prism for Windows (version 9.0.0, GraphPad Software, La Jolla, CA, USA) was used for statistical analysis of wet lab experiments. For genomics and transcriptomics data analyses, the Wilcoxon-rank sum test was used for comparison of 2 groups with continuous variables. The log-rank tests were used for comparing overall survival displayed as Kaplan–Meier survival curves. Differential expression analysis of transcripts was performed using the Wald test with DESeq2 v1.32.0 [75]. A gene list of differentially expressed genes was supplied to ReactomePA v1.44.0 [76] to identify enriched pathways with *p* values estimated by hypergeometric distribution. *p* values were adjusted for multiple testing using the Benjamini–Hochberg method and are indicated as adjusted *p* values. Genomics and transcriptomics data analyses were performed using the platform R v4.3.2 [78].

## Data Availability

All data needed to evaluate the conclusions in the paper are present in the paper and/or the Supplementary Materials. All RNA-seq data generated in this study have been deposited to the Gene Expression Omnibus (GEO) database with accession code X. WGS and RNA-seq data from metastatic bladder cancers were requested via the Hartwig Medical Foundation at https://www.hartwigmedicalfoundation.nl/en/data/data-access-request/, and approved under request number DR-314. The TCGA data for the bladder cancer cohort is publicly available at https://portal.gdc.cancer.gov./. Source data are provided with this paper. Additional data related to this paper may be requested from the authors.

---

**Supplemental Figure 1.**
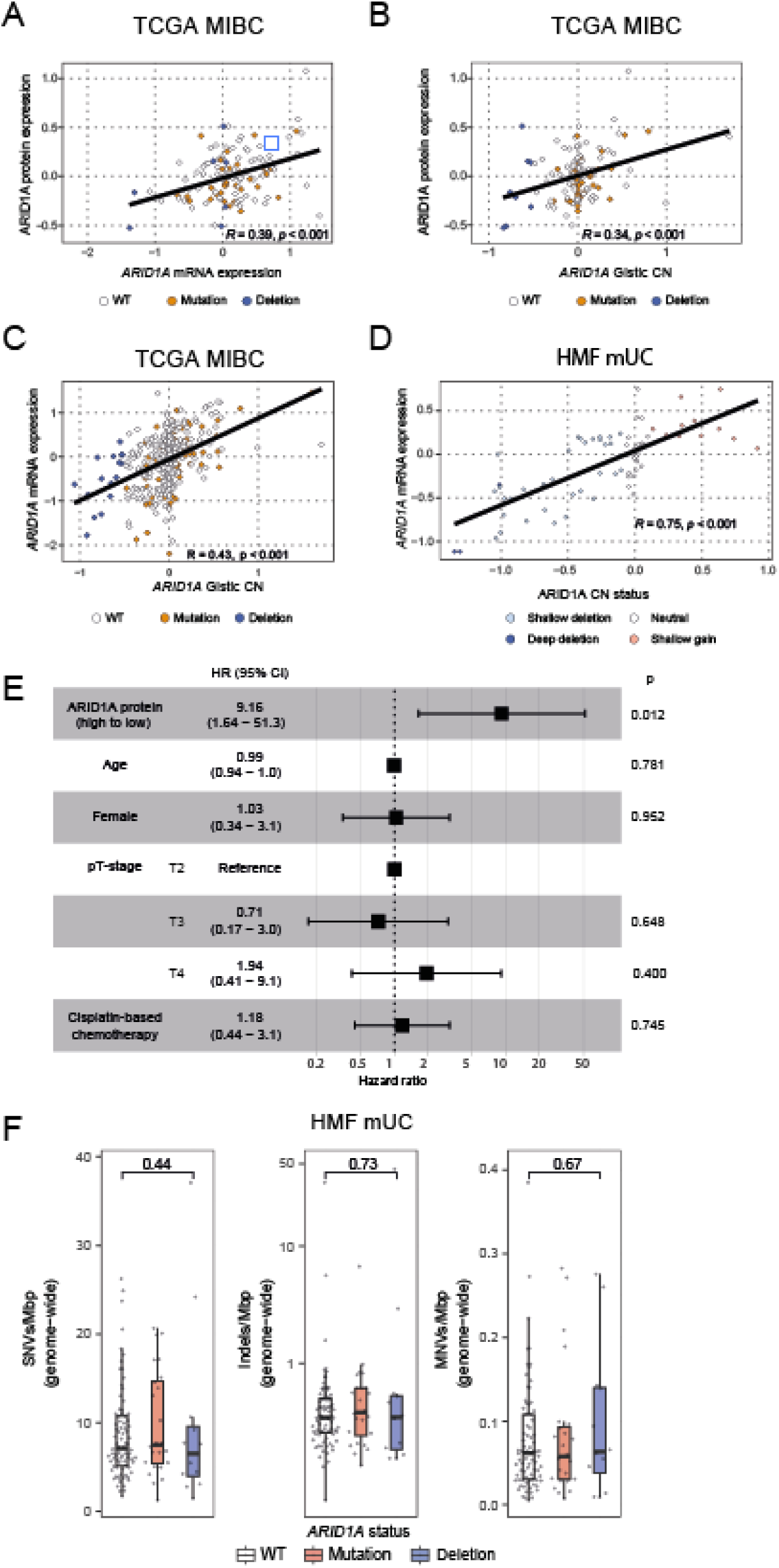
Pearson correlation between **A)** ARID1A protein expression levels and *ARID1A* mRNA expression, **B)** *ARID1A* protein expression and *ARID1A* gistic copy number levels, and between **C)** *ARID1A* mRNA expression and *ARID1A* gistic copy number levels in MIBC tumor samples*. **D)** Pearson correlation between *ARID1A* mRNA expression and *ARID1A* copy number status in mUC samples**. **E)** Overview of hazard ratios (HR) calculated for ARID1A protein expression and clinical features for MIBC patients*. Continuous variables were dichotomized based on the median and high vs. low is presented. Boxes indicate HR and horizontal lines show 95% confidence intervals (CI). **F)** Boxplots depicting the number of single-nucleotide variants (SNVs), insertions and deletions (indels), and multi-nucleotide variants (MNVs) per mega base pairs (MBp) in metastatic BC patients stratified on *ARID1A* mutation status. Two-sided Wilcoxon-rank sum test was applied to compare differences between samples with *ARID1A* WT and *ARID1A* deletions. For all graphs: WT = *ARID1A* wild type, Mutation = protein-coding mutation (excluding synonymous) and small insertions/deletions, Deletion = *ARID1A* deleted. *TCGA **HMF.

**Supplemental Figure 2.**
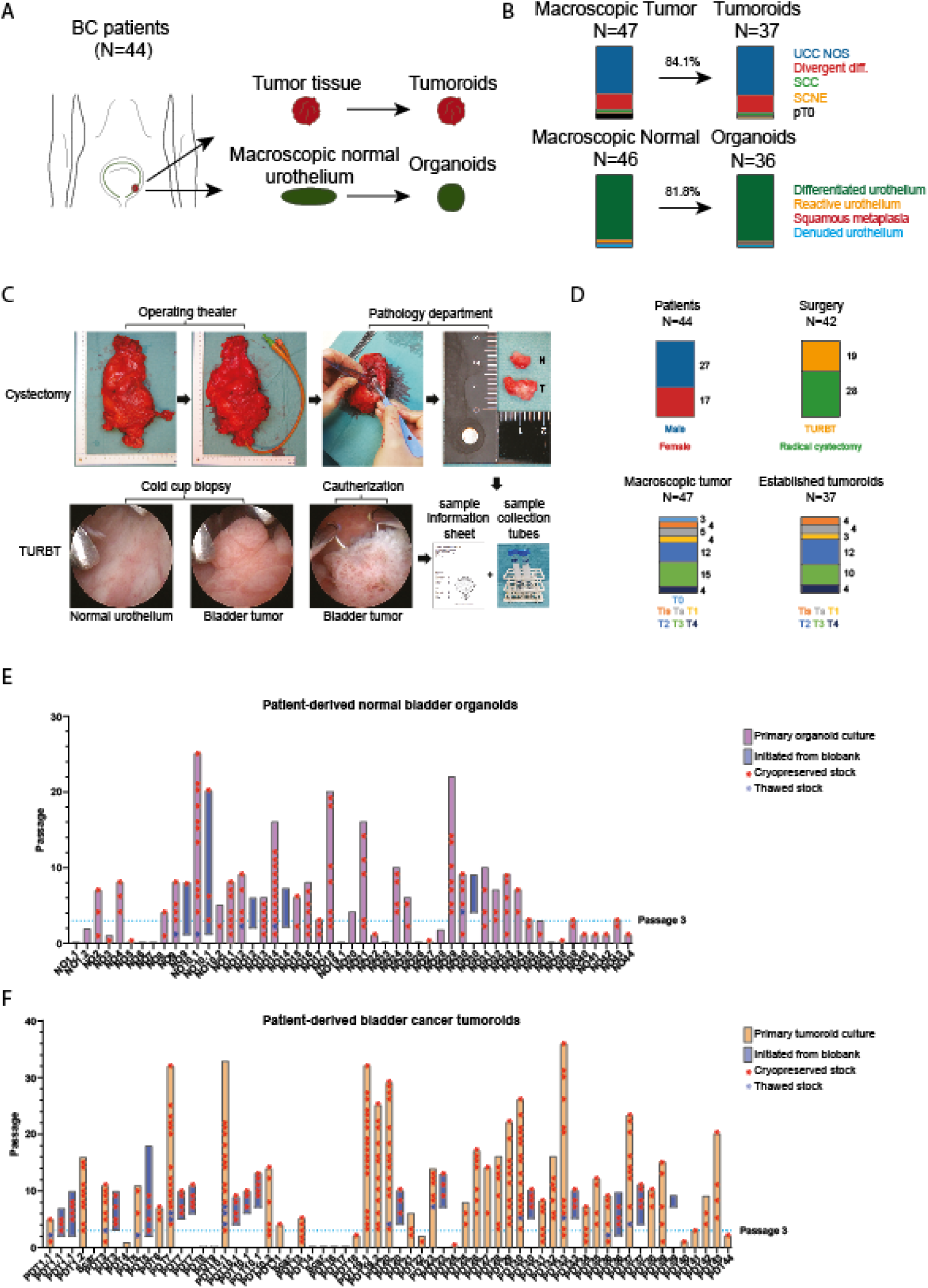
A) Diagram depicting the generation of patient-derived bladder tumoroids and normal organoids from primary tissues. Tumoroids are generated from tumor tissue, while organoids are generated from macroscopic normal urothelium. **B)** Top: Bar charts comparing the stratification of tumor samples with derived organoid lines based on histological subtype observed in the tumor samples UCC NOS = urothelial carcinoma not otherwise specified, Divergent diff. = divergent differentiation, SCC = squamous cell carcinoma, SCNE = small cell neuroendocrine. Bottom: Bar charts comparing stratification of macroscopic normal urothelium and derived organoid lines based on histological evaluation of macroscopic normal urothelium. **C)** Schematic depiction of sample acquisition. Bladders are instilled with cold preservation fluid immediately following radical cystectomy. Instillation occurs through an indwelling catheter, which is then plugged to prevent leakage during transport to the gross room. Bladders are kept inside an endobag and on ice to maintain sterility and preserve cell viability during transport. At the gross room, bladders are ventrally incised and normal (N) and tumor (T) samples are excised on a sterile field using disposable lancets and tweezers to prevent contamination. For transurethral resection of bladder tumor (TURBT), macroscopic normal urothelial tissue is sampled by cold cup biopsy prior to resection. Bladder tumor samples are thereafter acquired by cold cup biopsy or cautherization when biopsies were not possible due to limited visibility. Acquired samples are stored in collection tubes containing cold preservation fluid and are then transported to the laboratory facilities, along with a sample information sheet containing pseudomized baseline clinical information and study identification. **D)** Bar charts depicting patient characteristics and comparing stratification of all acquired macroscopic tumor tissue with that of all patient-derived tumoroid lines. N=39 patients underwent 42 surgical procedures. Top left: biological sex, top right: type of resection, bottom left: pathological tumor stage of all acquired samples, bottom right: pathological tumor stage corresponding to successfully initiated cultures. The composition of included patients and organoid lines is characteristic for a tertiary referral center and tumoroids were established from BC patients with various tumor stages without bias. **E)** Bar graph summarizing the established normal organoid biobank. Each bar represents one organoid line (purple bars) or derivative initiated from cryopreserved stocks (blue bars). Cryopreserved stocks are indicated by red asterisks, while thawed stocks are indicated by blue asterisks. **F)** Bar graph the established BC tumoroid biobank. Each bar represents one tumoroid line (yellow bars) or derivative initiated from cryopreserved stocks (blue bars). Cryopreserved stocks are indicated by red asterisks, while thawed stocks are indicated by blue asterisks.

**Supplemental Figure 3.**
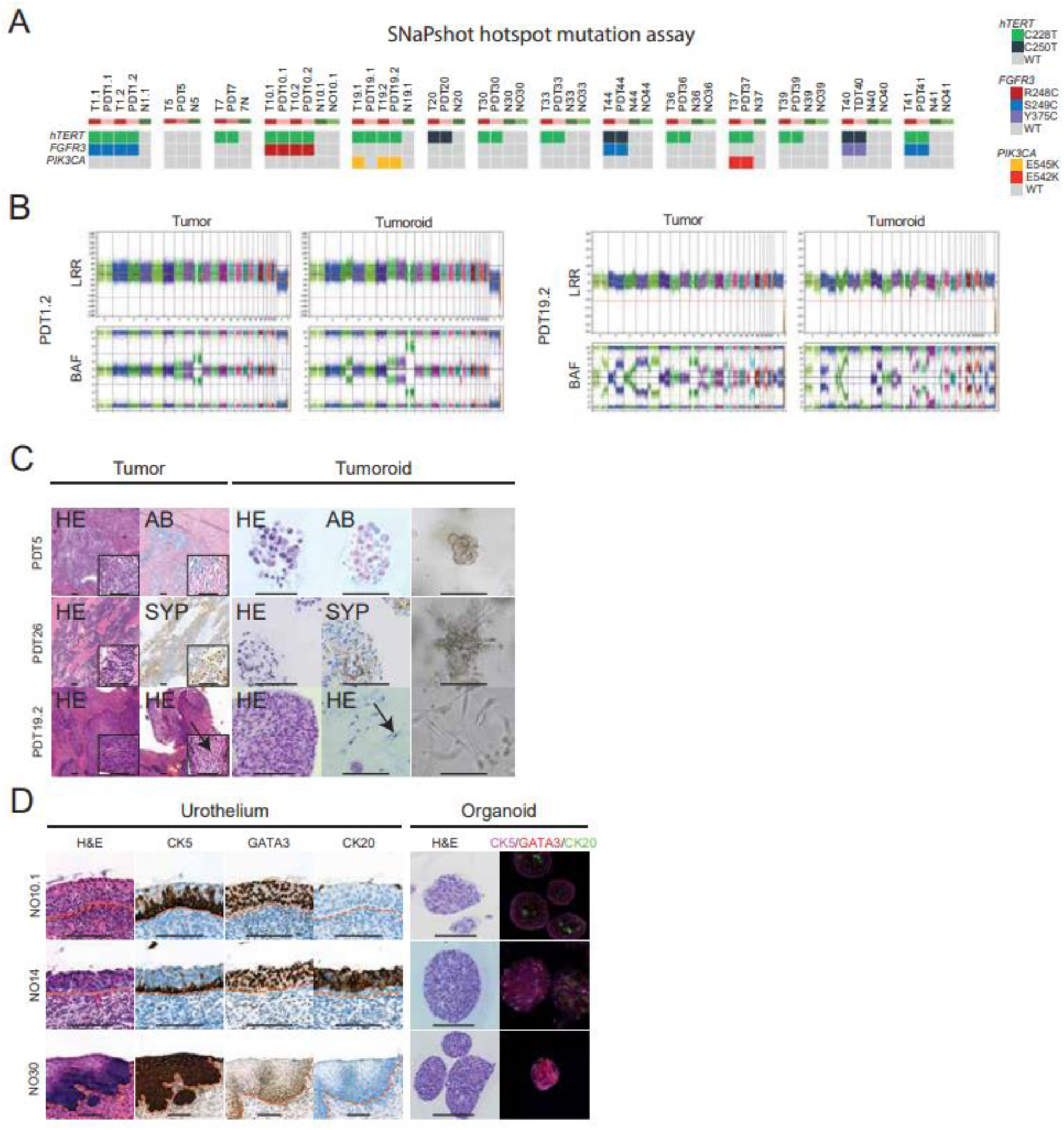
**A)** SNaPshot mutation analysis of patient tumor (T) and normal (N) samples and matched patient-derived tumoroid (PDT) and normal organoid (NO) cultures on recurrent somatic hotspot mutations in telomerase reverse transcriptase (hTERT), fibroblast growth factor receptor 3 (FGFR3), and Phosphatidylinositol-4,5-Bisphosphate 3-Kinase Catalytic Subunit Alpha (PIK3CA) genes, WT = wild type. **B)** Scatterplots illustrating genome wide copy number alterations depicted by Log R ratios (LRR) and B-allele frequency (BAF) from PDT1.2 and PDT19.2 tumor tissue and corresponding tumoroids. Note increased resolution of copy number alterations in tumoroid samples **C)** Comparative histological and immunohistochemical images of variant histology in bladder tumor tissue and corresponding tumoroid lines. Shown are representative examples of choroid differentiation, as well as small cell neuroendocrine and squamous cell carcinoma variant histology. Choroid differentiation (PDT5) shows signs of mucus production indicated by alcian blue (AB) positivity. Small cell neuro-endocrine bladder cancer (PDT26) stained positive for synaptophysin (SYP). Squamous cell carcinoma (PDT19.2) was identified by “tadpole” cells (black arrow) and keratinization. Two columns on the left demonstrate histological and (immune)histochemical images of bladder tumor tissue while the three columns on the right indicate patient-derived tumoroid lines. Scale bar = 50 μm. **D)** Histological evaluation of macroscopic normal bladder tissue and corresponding organoids. Two representative examples of normal urothelium (NO10.1 & NO14), in addition to one squamous metaplasia sample (NO31) are shown. Organoids and originating tissue were compared by H&E staining, while expression of urothelial differentiation markers was investigated by IHC (original tissue) and IF (organoids) as indicated. (scale bar = 50 μm).

**Supplemental Figure 4.**
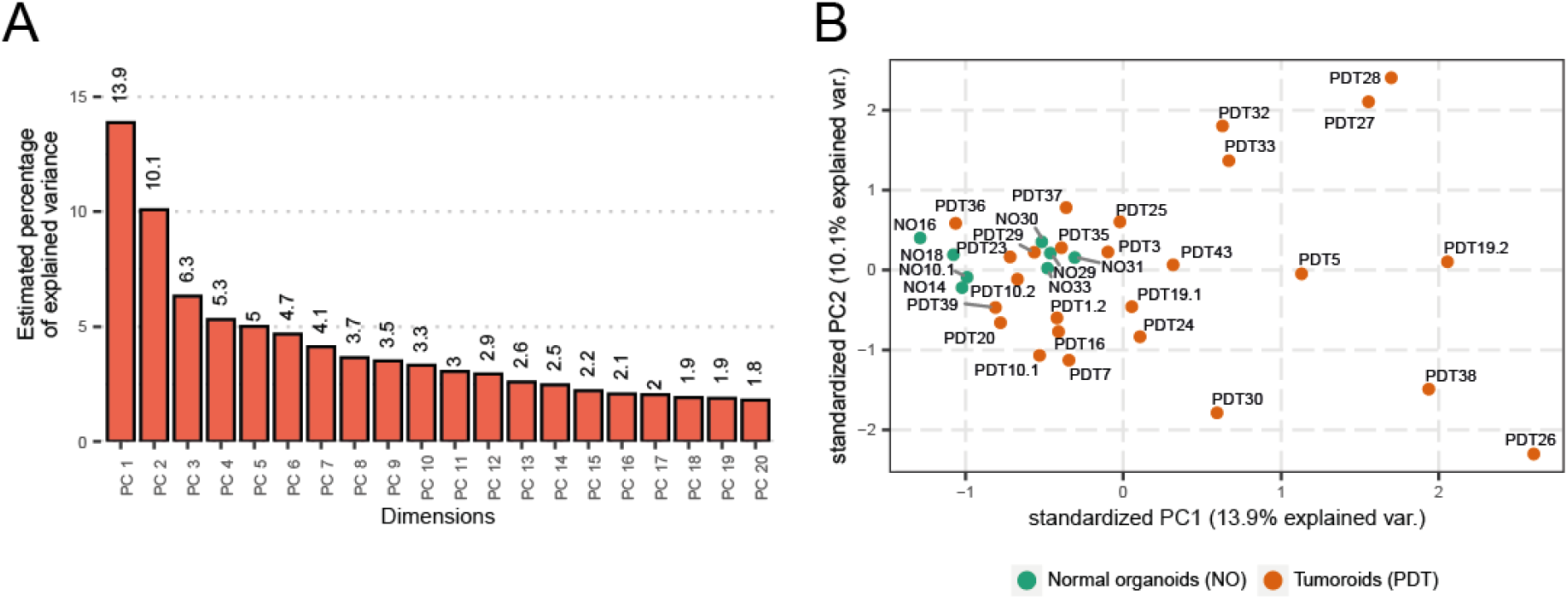
**A)** Bar graph depicting estimated variance in gene-expression by the top 20 principal components. **B)** Distribution of tumoroids and normal bladder organoids within the first two principal components (PCA1-2).

**Supplemental Figure 5.**
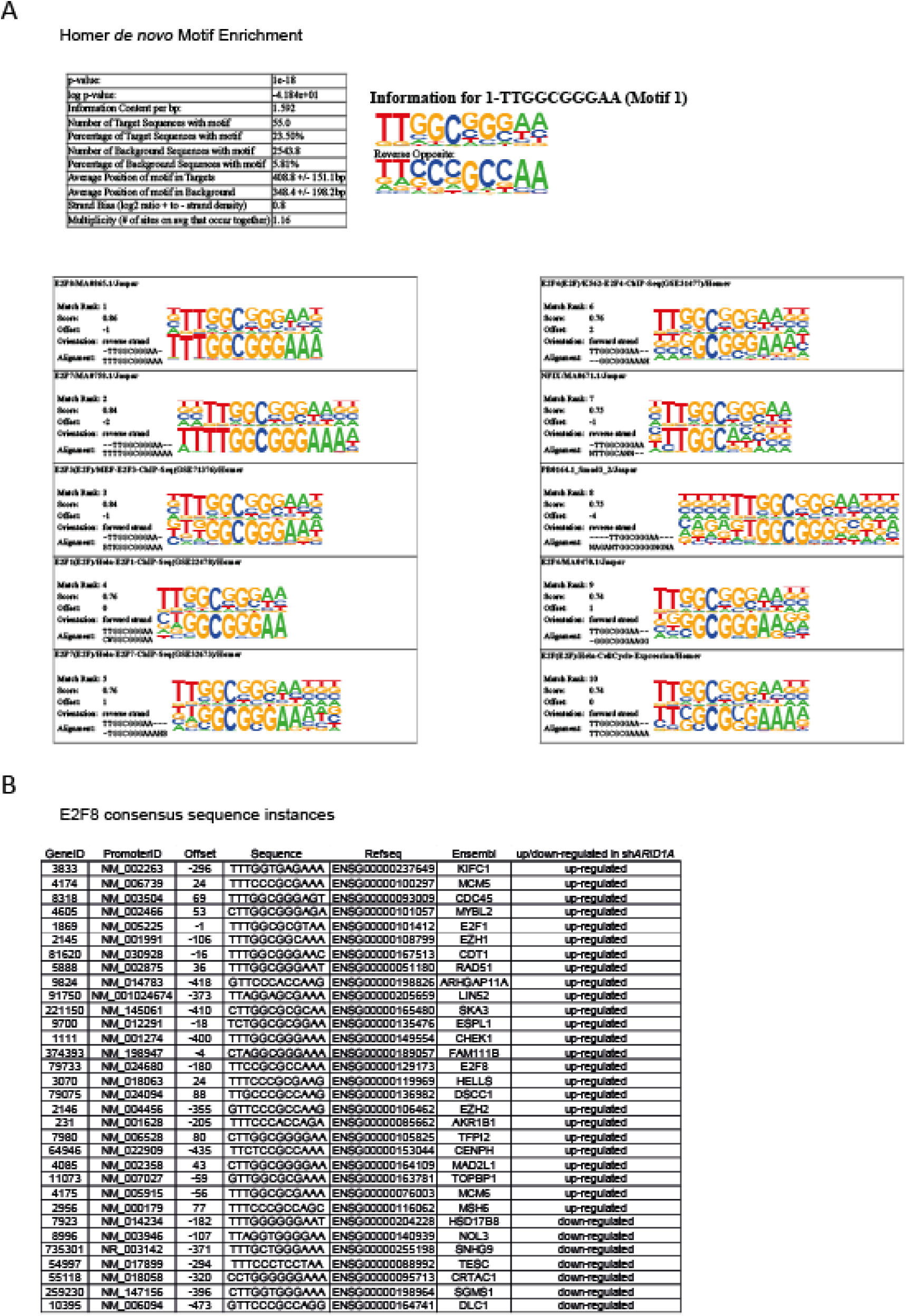
**A)** *De novo* binding motif (Motif 1) and matched known binding sequences as predicted by Homer. **B)** E2F8 consensus sequence instances in dysregulated genes upon ARID1A knock-down in bladder organoids.

**Supplemental Figure 6.**
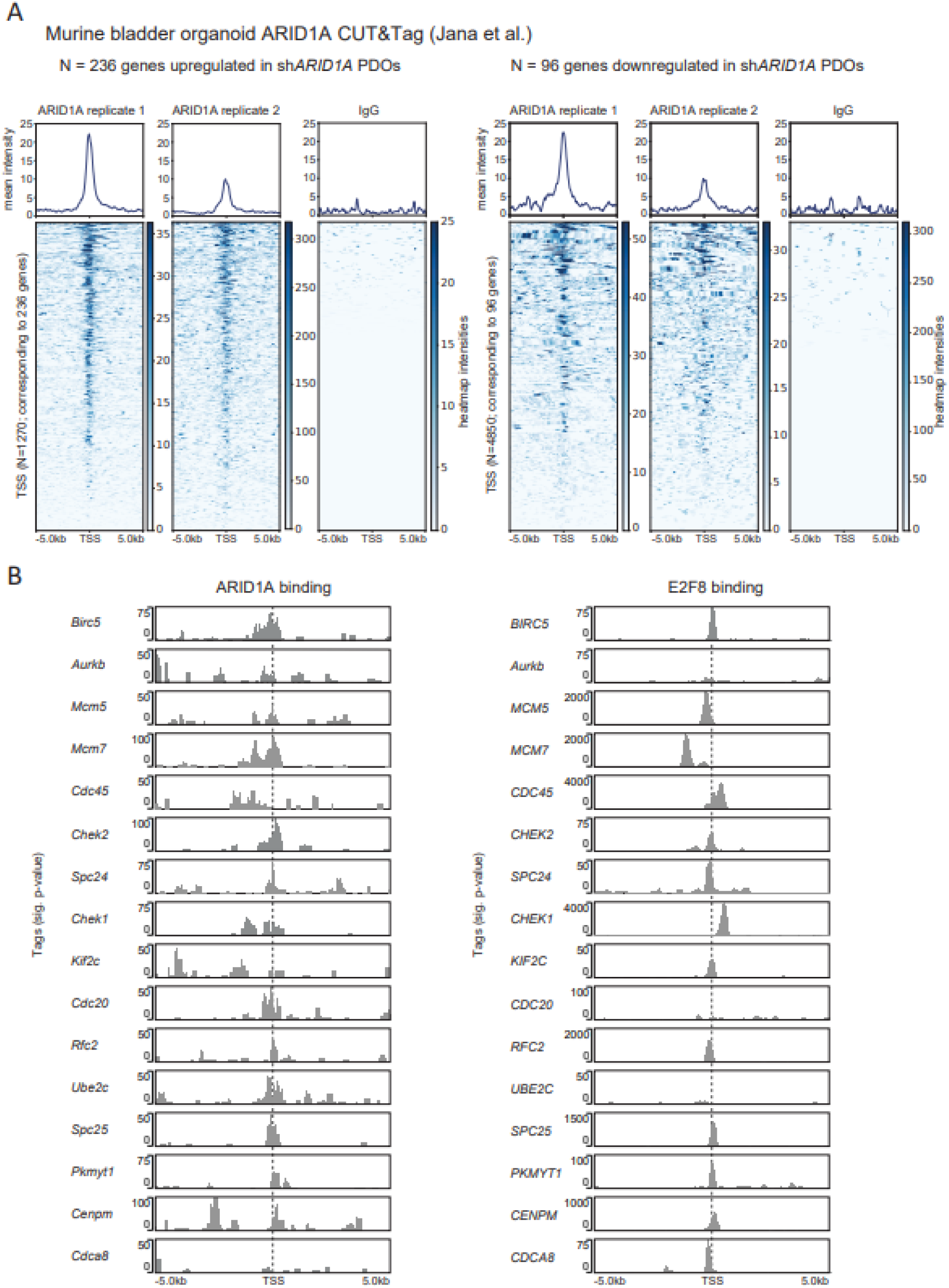
**A)** Histograms (top) and heatmaps (bottom) depicting mean ARID1A binding intensities around 1270 transcription start sites (TSS) corresponding to N = 236 genes upregulated (left) or N=96 downregulated (right) upon ARID1A knock-down in normal organoids. N= 10 genes did not have a murine orthologue and where excluded from analysis. CUT&Tag analysis was performed in murine urothelial organoid cells with ARID1A-directed antibodies, using IgG as control as indicated. Data was repurposed from Jana et al. **B)** Left: ARID1A occupancy at transcription start sites of 16 candidate genes in murine urothelial organoids. Data was repurposed from Jana et al. (29). Right: E2F8 occupancy at transcription start sites of 16 candidate genes in K562 myeloid progenitor cells. Data was repurposed from the ENCODE project (35).

**Supplemental Figure 7.**
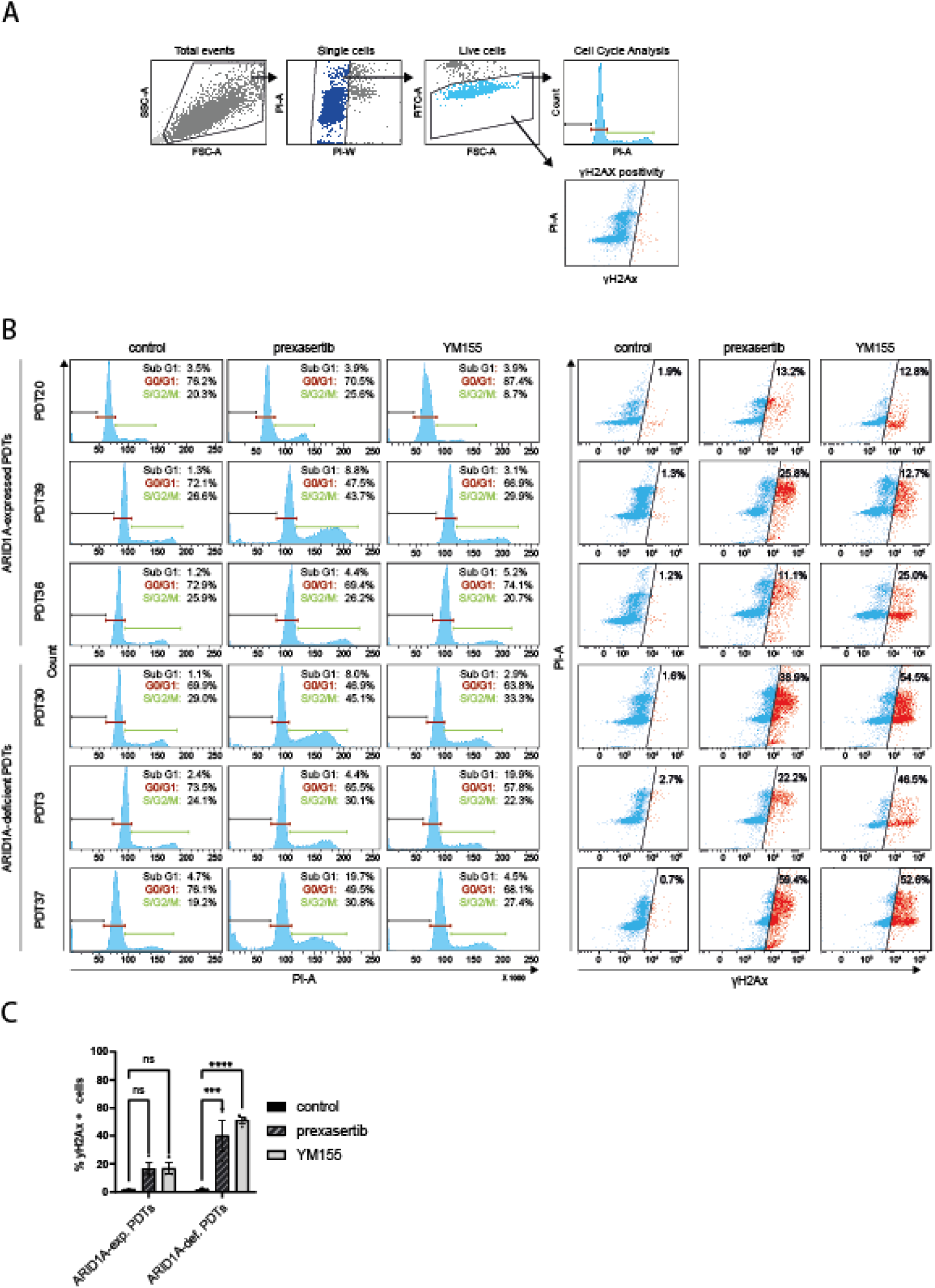
**A)** Sequential gating strategy to identify single cells: initial gating on FSC-A vs SSCA to define the region of interest by excluding very small particles and multicellular aggregates, on the left, followed by single cells gating in PI-A vs PI-W, cell cycle analysis using a mono-dimensional PI-A histogram in a linear scale, and gating strategy to define the percentage of γH2AX+ cells in the live fraction. **B)** Flow cytometry on ARID1A-expressing tumoroids (PDT20, PDT36, PDT39) and ARID1A-deficient tumoroids (PDT3, PDT30, and PDT37) treated with 32nM prexasertib or 50nM YM155 for two days. Gating on single live cells was followed by cell cycle analysis (left) and measurement of the fraction of cells with phosphorylated H2Ax (γH2Ax) (right). **C)** Bar graph depicting γH2Ax+ fraction of ARID1A-expressing (PDT20, PDT36, PDT39) and ARID1A-deficient (PDT3, PDT30, and PDT37) tumoroids treated with 32nM prexasertib or 50nM YM155 compared to untreated control. Gating on single live cells was followed by measurement of γH2Ax+ positive fraction, as shown in B. Drug treated ARID1A-expressing and ARID1A-deficient tumoroids are compared to the respective untreated controls, however, source data corresponds to that presented in Figure 5C. Data are represented as mean ± SEM.

**Supplemental Figure 8.**
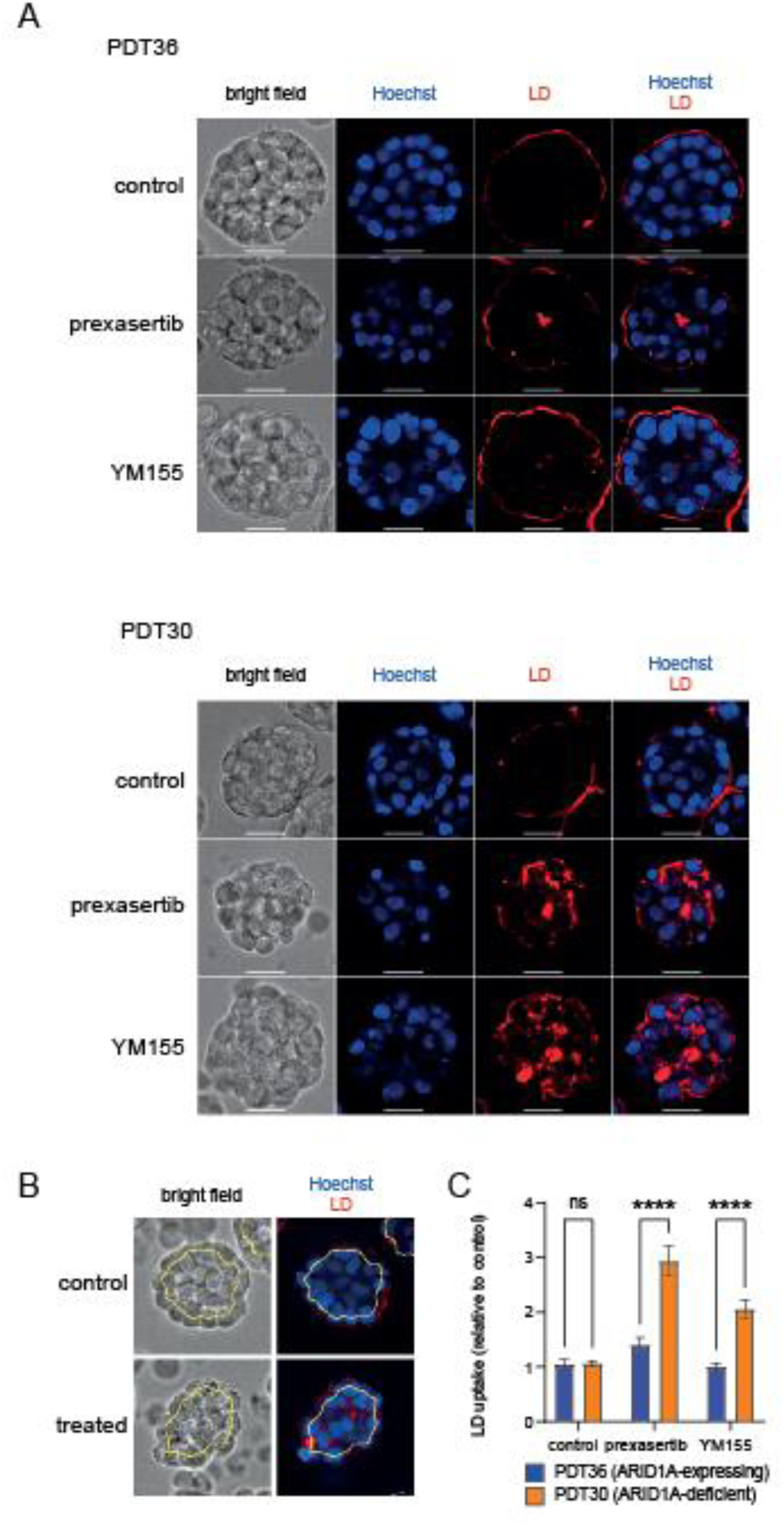
**A)** Representative brightfield and fluorescence staining showing fixable Live/Dead (LD) cell staining (red) in *ARID1A*-expressing (PDT36) and *ARID1A*-deficient (PDT30) bladder tumoroids treated for 48 hours with 32nM prexasertib or 50nM YM155, compared to untreated controls. Hoechst (blue) was used as a counter staining. Raw images were exported from Harmony software and brightness and contrast was adjusted in imageJ **B)** Representative images depicting automated tumoroid segmentation (yellow line) by the Harmony software to select the inner 60% of tumoroid surface area in order to quantify tumoroid uptake of fixable live/dead staining. Processed Images were exported from Harmony software and contain gamma correction as implemented by Harmony software. Segmentation line was manually accentuated in illustrator **C)** Bar graph depicting mean uptake of fixable LD staining per tumoroid. LD uptake was measured in the inner 60% of tumoroid surface area. At least 100 tumoroids were measured per condition and no gamma correction was applied prior to measurement. Data are presented as mean +-SEM. **P<0.005, ****P<0.00005.

**Supplemental Figure 9.**
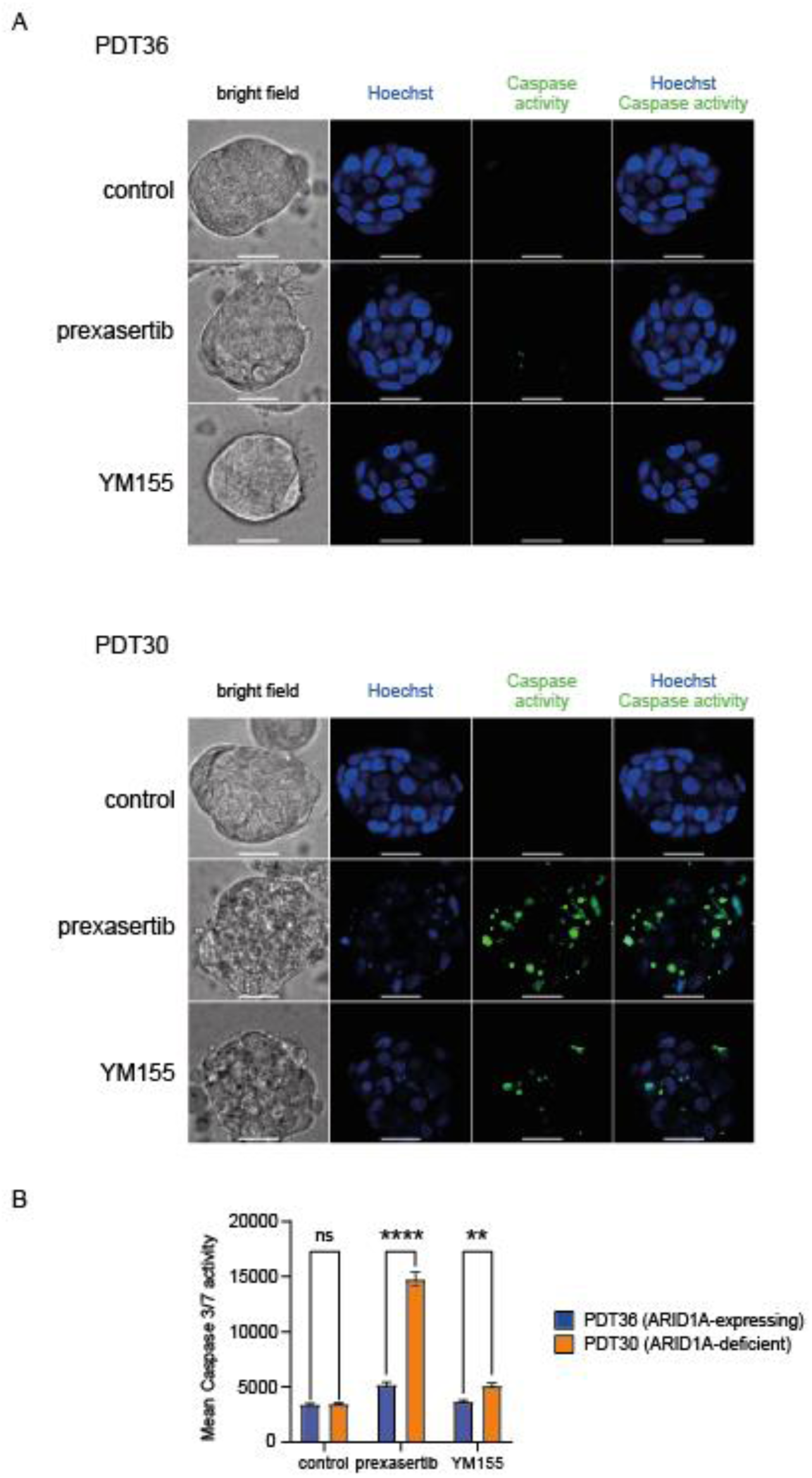
**A)** Representative bright field and fluorescence staining showing caspase 3/7 activity (green) in *ARID1A*-expressing (PDT36) and *ARID1A*-deficient (PDT30) bladder tumoroids treated for 48 hours with 32nM prexasertib or 50nM YM155, compared to untreated controls. Hoechst (blue) was used as a counter staining. **B)** Bar graph depicting mean staining intensity (Caspase 3/7 activity) per tumoroid. At least 49 tumoroids were measured per condition and data are presented as mean +-SEM. **P<0.005, ****P<0.00005.

**Supplemental Figure 10.**
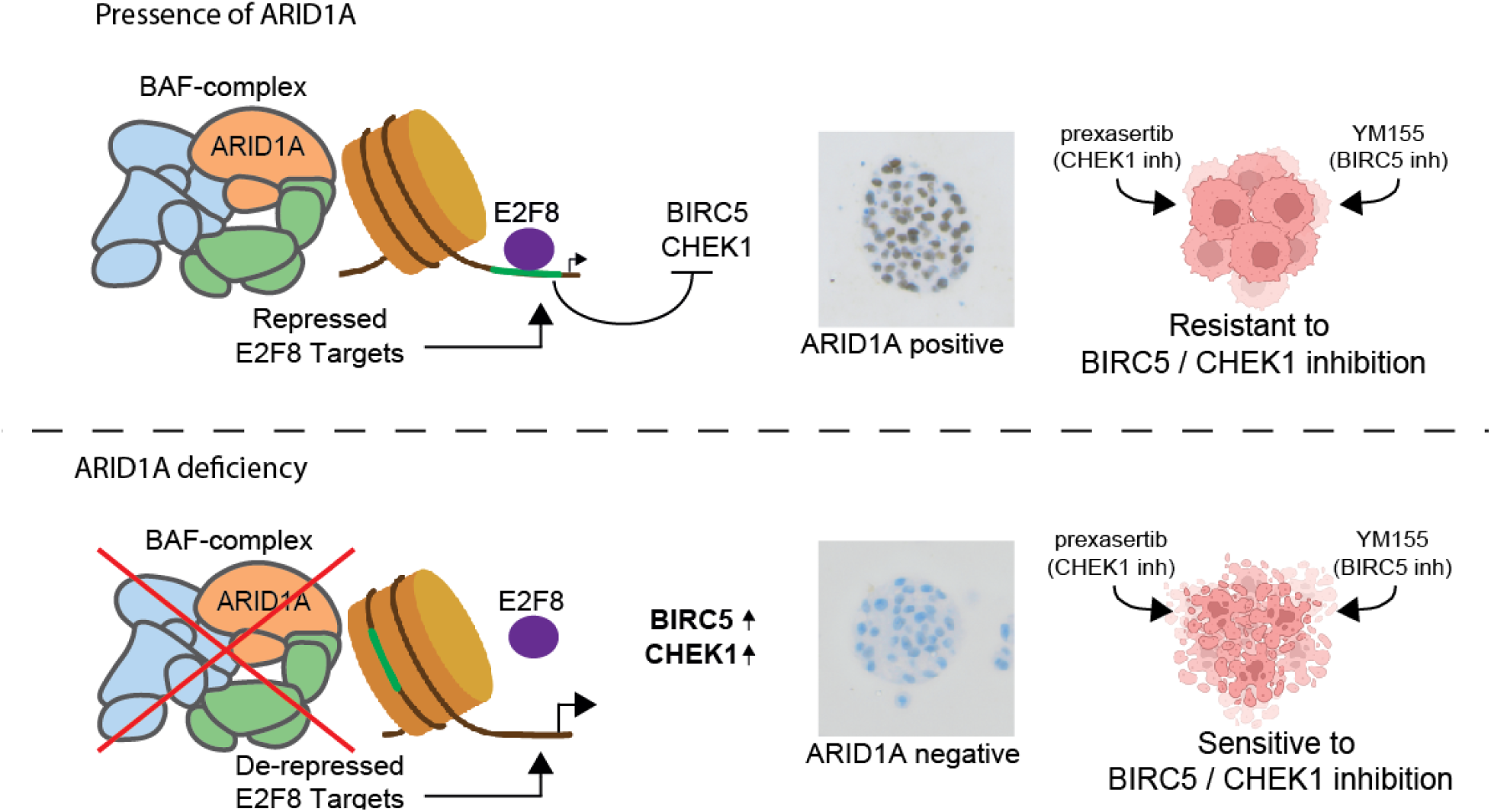
Theoretical model for the interaction of E2F8 and ARID1A and regulation of CHEK1 and BIRC5 expression in ARID1A-expressing and ARID1A-deficient cells. **Top:** ARID1A and the BAF complex reposition the nucleosome, exposing E2F8 binding motifs. Upon binding, E2F8 represses its target genes, including CHEK1 and BIRC5. As a result, CHEK1 and BIRC5 expression is generally repressed, making ARID1A-expressing cells relatively resistant to BIRC5 and CHEK1 inhibition. **Bottom:** Loss of ARID1A expression disrupts the BAF complex’s ability to reposition nucleosomes, hindering E2F8 binding and leading to de-repression of BIRC5 and CHECK1. In turn, upregulated expression of BIRC5 and CHEK1 can be therapeutically targeted by small molecule inhibitors, thereby selectively eliminating *ARID1A*-deficient BC cells.

